# The substrate changes matrix metalloprotease-1 dynamics, allostery, and ligand binding affinity

**DOI:** 10.1101/2025.02.08.637235

**Authors:** Chase Harms, Anthony Nash, Susanta K. Sarkar

## Abstract

Using single-molecule experiments, ensemble assays, and Molecular Dynamics (MD) simulations, we showed that collagen actively influences MMP1 interdomain dynamics. This paper extends the previous experimental work to validate MD simulations and gain more insights into MMP1-collagen interactions. We include virtual screening with free energy calculations to uncover key insights regarding substrate-dependent dynamics, allostery, and ligand binding. Our findings reveal that MMP1 interdomain dynamics are influenced by the strain in collagen, and collagen-specific allosteric residues exhibit strong correlations with the active site. Additionally, allosteric paths connecting these residues to the active site bypass the linker region. Collagen also induces changes in the configurations of the catalytic motif. Furthermore, MD simulation and virtual screening identify molecules that bind to allosteric residues, alter MMP1 dynamics, and potentially change activity. Free energy calculations highlight the essential role of the substrate in small molecule screening against a protein. We have developed a comprehensive framework for identifying substrate-specific allosteric residues in MMP1 and screening small molecules incorporating protein dynamics in the presence of substrates. These results significantly impact our understanding of MMP1 interactions with collagen fibrils and pave the way for selectively modulating specific MMP1 functions without altering its other functions in future applications.

**Significance:** In this paper, we validated all-atom simulations of free and collagen-bound MMP1 and gained insights into MMP1 function. We have found that collagen changes dynamics, allostery, and ligand binding in MMP1. Collagen strain, known to be age-dependent, also changes MMP1 dynamics and ligand binding. Interdomain allosteric paths do not go through the linker connecting the MMP1 domains. Instead, paths from the collagen-specific allosteric residue to the active site have non-bonded interactions. Screening of small molecules against collagen-specific MMP1 residue suggests that collagen alters ligand binding to MMP1. These results will significantly impact understanding MMPs and small molecule screening against proteins. Substrate-specific allosteric residues may enable controlling one function of MMPs without altering their other functions, leading to fewer side effects.

## Introduction

MMP1, a collagenase in the 23-member MMP family, can degrade type-1 collagen’s triple-helical structure (1). Type-1 collagen, one of the 28 human collagen types (2), is the most abundant collagen type and the sixth most abundant protein type in the body (3). Collagen fibrils provide a scaffold for cellular organization and tissue integrity, and its degradation by MMP1 is still unclear (4). The degradation requires both the catalytic and hemopexin domains, suggesting allosteric communications (5). Experimentally, a mixture of the two MMP1 domains purified separately can degrade triple-helical collagen (5) without the physical linker, suggesting allosteric communications via substrate (6). Recently, we showed that the distributions and time-correlations of MMP1 conformations on type-1 collagen correlate with activity. Ligands such as tetracycline have known effects on the function and can also modulate the MMP1 dynamics (6). Also, interdomain dynamics are substrate-dependent and show significant differences with different substrates, including type-1 collagen fibril (6), fibrin (7), alpha-synuclein (aSyn) (8), and amyloid-beta (Aβ) aggregates (9).

To understand how MMP1 adapts to different substrates, we note that the catalytic domains of MMPs have mostly conserved sequences despite their diversity of substrates and activities. However, the hemopexin domains have significant variations in sequence (10, 11). These observations suggest that interdomain allosteric communications between the two domains may be critical in MMPs’ substrate specificity and enzymatic activity.

Allosteric communication occurs when a signal transfers from one site to a distant location where the protein functions, such as catalysis, occur. There are three main paradigms of allostery: induced fit (12-14), conformational selection (15-24), and allosteric networks (25-28). The “induced fit” model suggests that a protein undergoes a stepwise change in conformation upon interaction with a ligand for mutual best fit (Koshland-Nemethy-Filmer or KNF model) (12). In contrast, the “conformational selection” model by Monod, Wyman, and Changeux (MWC) suggests that allosteric proteins are oligomers of identical subunits. Each subunit can exist in at least two (relaxed and tensed) conformations in equilibrium. The ligand can bind to both conformations and cooperatively force all subunits to assume the same conformation but have a higher affinity for the relaxed state, shifting the equilibrium toward the relaxed state (29). Allosteric communications can occur even when the average state does not change upon ligand binding (30). Nussinov’s group extended the MWC model from two states to ensembles of multiple states and suggested that the ligand preferentially interacts with one of many protein conformations and shifts the equilibrium toward the selected conformation (18, 19, 22, 31). Finally, Ranganathan’s group showed that evolutionarily conserved amino acid networks mediate allosteric communication (27, 32). Nussinov’s group suggested that all proteins are allosteric (33) and provided a unified view of allostery (34). Researchers have explained allostery from other perspectives, such as Le Chatelier’s principle (35), a network of tertiary and quaternary motion (36), perturbations (37), information transfer (38), and conformational dynamics (39, 40). However, it is unclear how substrates can influence allosteric communications when the binding affinity is strong.

This paper shows that collagen induces changes in MMP1 dynamics and the catalytic motif configuration. Building on our previous publication on single-molecule measurements of MMP1 dynamics, we performed all-atom MD simulations of MMP1 bound to a triple-helical collagen model (PDB ID 4AUO) (41). The results revealed that MMP1 dynamics depend on collagen strain, and we identified residues in the hemopexin domain, defined as the allosteric residues or “allosteric fingerprint,” with strong correlations with the catalytic motif residues in the active site. Some allosteric residues in free MMP1 lose correlations, and some residues develop correlations with the catalytic motif upon collagen binding. As a proof of principle, we performed a virtual screening of small molecules against MMP1. We demonstrated how changes to the collagen substrate affect small molecule binding free energy to the proposed allosteric binding site. Identification of collagen-specific allosteric residues enables more specific screening of small molecules. In the future, identifying substrate-specific allosteric residues or “allosteric fingerprints” may allow for controlling one MMP1 function without affecting its other functions.

## Results and discussion

We started with PDB ID 4AUO (41) and constructed six models of MMP1 in its active (E219) and catalytically inactive point mutant (E219 to Q219) forms for six models. For active and inactive MMP1, we modeled (a) free MMP1, in which no collagen is present; (b) collagen-bound MMP1 with unrestrained collagen; and (c) collagen-bound MMP1 with 1000 kJ/mol position restraints applied to the alpha-carbon atoms of the collagen. The three helical chains making the collagen molecule were periodic, crossing the periodic boundary condition and forming an uninterrupted chain to mimic the physical properties of a full-length collagen molecule. **Figure 1** shows two representative models. For details of model constructions, see methods.

**Figure 1.**
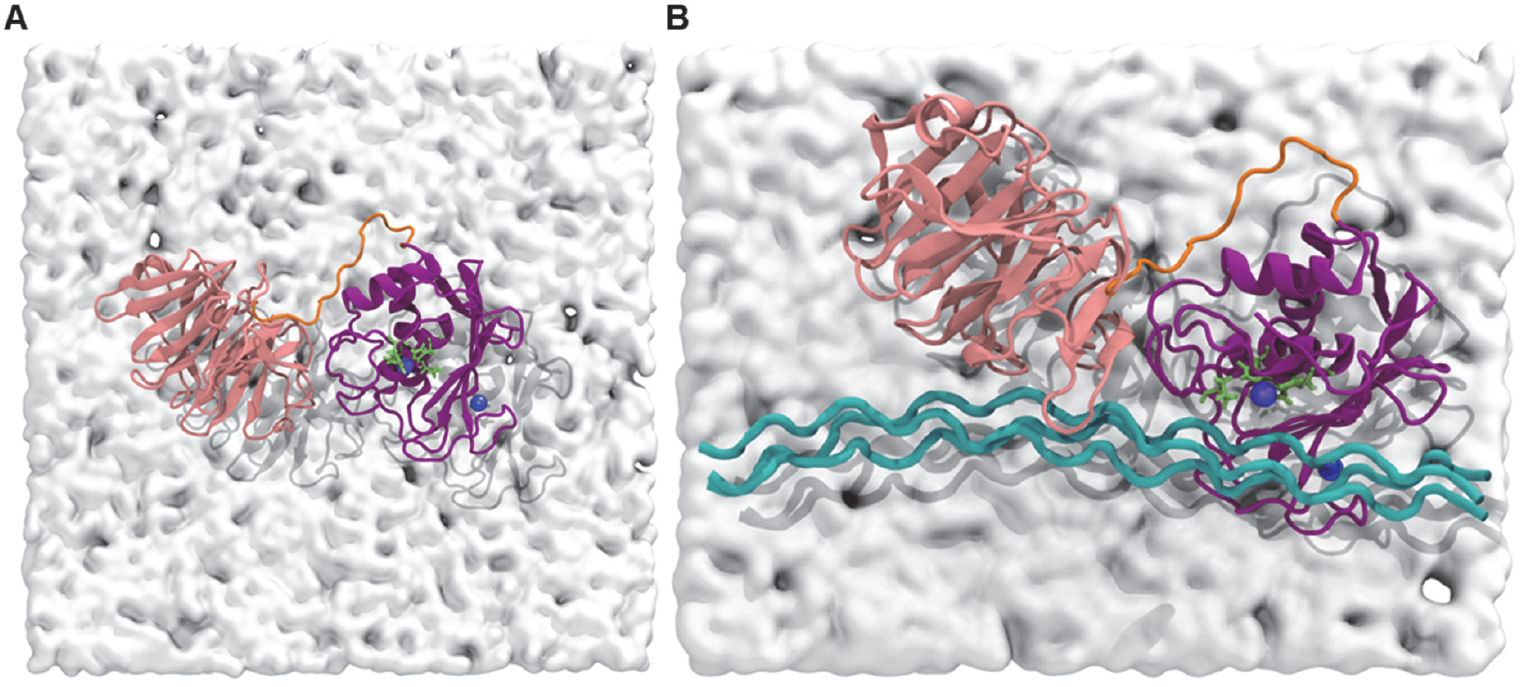
Representative models of (A) free MMP1 and (B) collagen-bound MMP1. The catalytic domain (F100–Y260) is purple, the linker residues (G261–C278) are orange, and the hemopexin domain (D279–C466) is pink. The catalytic and structural zinc ions are in blue; the three histidine residues that coordinate the catalytic zinc and the active site glutamic acid (E219) are in green. The collagen molecule was made periodic across the unit cell, and a layer of water residues was included to provide a visual perspective on the difference in unit cell size between the two models.

We simulated all-atom dynamics of free and collagen-bound MMP1, solvated in water, using an integration time step of two femtoseconds. We recorded coordinates every five picoseconds (time resolution) unless otherwise stated. All simulations were subjected to a thorough gradient descent energy minimization process until all atoms settled below the maximum force threshold of 500 kJ^-1^ mol^-1^ nm^-1^, followed by a series of relaxation steps. First, 1000 kJ/mol position restraints were applied to all heavy atoms of MMP1 and collagen. An NVT simulation, where the number of atoms N, volume V, and temperature T are fixed, was performed for 50 ps using the V-rescale thermostat. Velocities from the final NVT simulation step were applied to start a 50 ps NPT simulation, where the number of atoms N, pressure P, and temperature T are fixed, using a C-rescale pressure coupling. After 50 ps of the NPT simulation, we reduced the position restraints to 100 kJ/mol and simulated an additional one nanosecond. Unless otherwise stated, all position restraints were removed, and the coordinates and velocities from the last timestep were used to begin production runs. We checked the system Root-Mean-Squared-Deviation (RMSD), temperature, and pressure for all simulations (as an example, see the supplementary **Figure S1** for collagen-bound MMP1 at 1000 kJ/mol). For details, see methods and supplementary information. We compared with experimental dynamics in our previous publication to validate simulations, extracted relevant data and analyzed them as presented below.

### MMP1 conformation changes upon collagen binding and depends on the collagen strain

Previously, we showed that MMP1 activity on collagen fibril degradation correlates with interdomain dynamics, and an open conformation where the catalytic and hemopexin domains are well-separated is relevant for catalytic activity (6). Literature shows that the collagen strain alters the enzymatic degradation of fibrils (42); specifically, lower strains inhibit degradation (43-46), whereas higher strains increase degradation (47-49). Some studies have shown that as strain varies from low to high, inhibition of degradation changes to enhancement at a threshold strain (50-52). Since collagen strain varies with age and affects collagen degradation in the human body (53), we investigated the molecular mechanism behind strain-dependent collagen degradation. Our previous papers (6-8) and others (49) have shown that MMP1 dynamics play an essential role in MMP1 activity.

To gain a deeper understanding of strain-dependent dynamics and activity, we compared experimentally measured MMP1 dynamics at the single molecule level in our previous paper (6) using all-atom molecular dynamics simulations (**Figure 2**). We simulated the dynamics of each of the three models in their inactive and active form for 100 ns. The histograms of domain separation fit a sum of two Gaussians, similar to the experimental histograms reported in a previous publication (6). For active MMP1, the interdomain separation, quantified by *mean* ± *standard deviation*, is 3.36 ± 0.03 nm for free (**Figure 2A**), 4.21 ± 0.16 nm upon binding collagen without restraint (**Figure 2B**), and 4.31 ± 0.11 nm with 1000 kJ/mol restraint on collagen (**Figure 2C**), respectively. In contrast, the interdomain separations of inactive mutant (E219Q) of MMP1 are 3.38 ± 0.04 nm for free, 4.28 ± 0.16 nm upon binding collagen without restraint, and 4.35 ± 0.11 nm with 1000 kJ/mol restraint on collagen, respectively. **Figures 2A-C** suggest an increase in collagen strain increases the interdomain separation of MMP1. This is consistent with our previous publication, where we showed that increased MMP1 activity is correlated with increased interdomain separation (6). Taken together with publications from other researchers that higher strains increase collagen degradation (47-49), we posit that the interdomain distance of MMP1 can be used to predict MMP1 activity.

**Figure 2.**
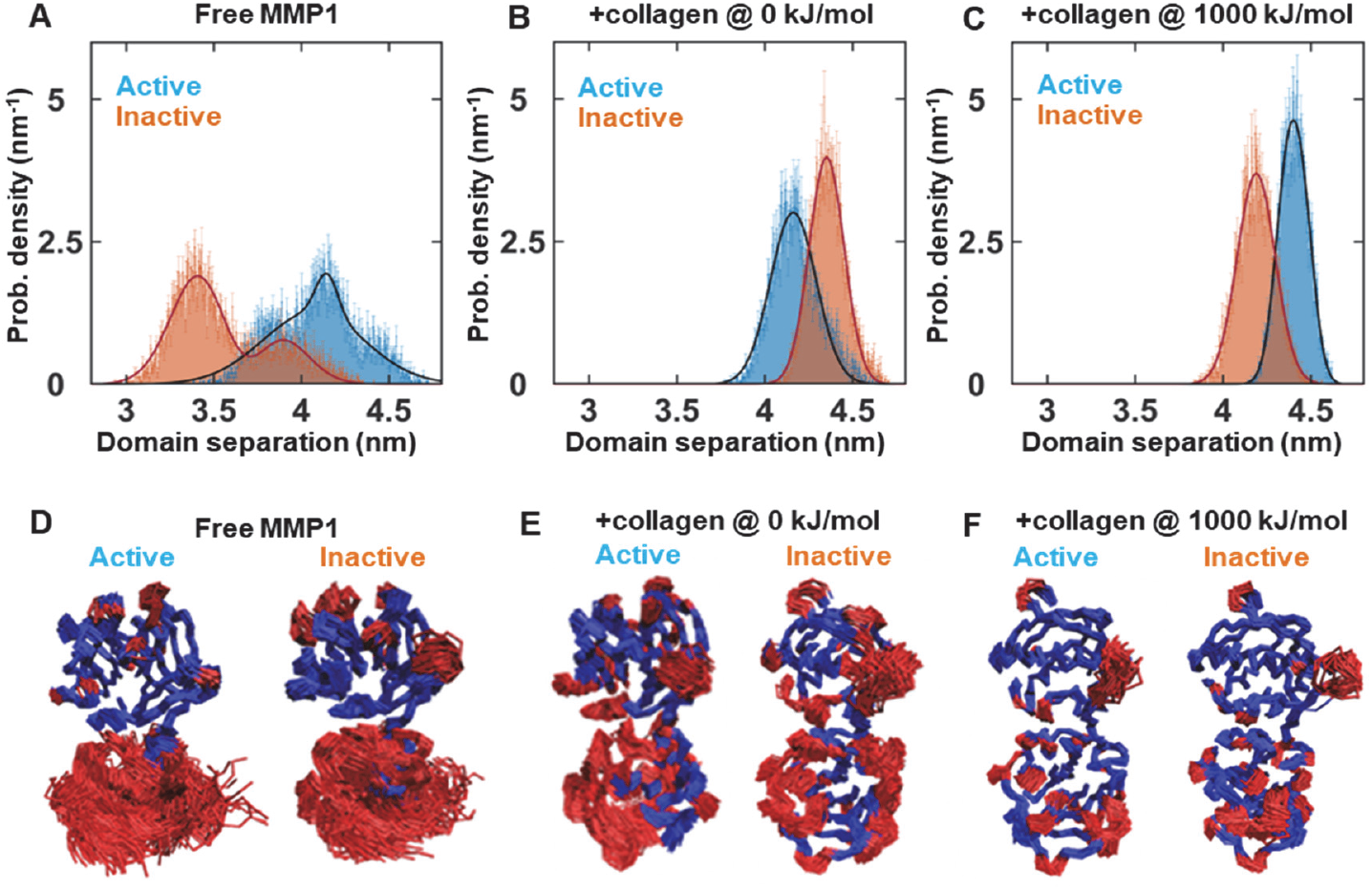
MMP1 dynamics depend on collagen strain. Area-normalized histograms of MMP1 interdomain distance for active (blue) and inactive (orange) MMP1: without collagen (**A**); bound to collagen without restraints (**B**); and with 1,000 kJ/mol restraints on the collagen alpha carbons (**C**). Error bars in histograms represent the square roots of histogram bin counts. Solid lines represent the best fit to a sum of two Gaussians. We superimposed 20,000 structures of MMP1 from a 100 ns-long simulation with 5 ps time resolution and 2 fs integration time to show how domains fluctuate in active and inactive forms for (**D**) free MMP1, (**E**) collagen-bound MMP1 without position restraints on the collagen-alpha-carbon-atoms, and (**F**) collagen-bound MMP1 with 1000 kJ/mol position restraints on the collagen alpha carbon atoms. The catalytic domain (bottom domain) has more flexibility than the hemopexin domain (top domain). Blue represents the least mobile residues (see methods for details). Collagen reduces MMP1 flexibility significantly. This is consistent with individual residues’ Root-Mean-Squared-Fluctuations (RMSF) (**Figure S2**).

We quantified fluctuations in different parts of MMP1 to determine how collagen alters those dynamics (**Figures 2D-F**). The difference between active and inactive forms was subtle, although the fluctuations were significantly higher in the catalytic domain than in the hemopexin domain in both cases. Interestingly, introducing the collagen substrate reduced flexibility in MMP1, even more so when the strain was introduced to the collagen alpha carbons. Additionally, we calculated the RMSF of the alpha-carbon atoms in MMP1 (**Figure S2**), which also revealed the higher fluctuations in the catalytic domain residues.

The observations of strain-dependent MMP1 dynamics and corresponding variations in the interdomain separation have several implications. First, we showed that active MMP1 prefers larger interdomain separation using single-molecule Forster Resonance Energy Transfer measurements (6). A comparison with our simulations suggests that experimental observations match only with restraints on the collagen alpha carbons. The agreement with restrained collagen is plausible because strain is introduced as collagen monomers self-assemble into fibrils (54). Second, the strain-dependent dynamics provide new insights into the debate about closed and open MMP1 conformations’ functional relevance. Prior studies supported that a larger separation between the two MMP1 domains (open conformation) is necessary for function, based on Nuclear Magnetic Resonance, small-angle X-ray Scattering, and MD simulation studies (55-58). In contrast, X-ray crystallography (PDB ID 4AUO) supports that the two domains are closer (closed conformation) for activity (41). Finally, single-molecule measurements (6), simulations presented in this paper, and reports by other researchers (55, 56, 59) support that MMP1 exists in an equilibrium between open and closed conformations. However, the active MMP1 mainly adopts the open conformation on collagen, and the preference increases as the strain in collagen increases. With these validations of simulations using published results (6), we continued our analyses using only the active form of MMP1 to gain insights into allostery and ligand binding.

### There are strong allosteric communications between the hemopexin and catalytic domain

We used simulations in **Figure 2** to identify allosteric residues exhibiting correlations with the residues of the catalytic motif and define the changes at the active site. We calculated correlations of fluctuations, as described in previous publications (6-8). We divided the simulated trajectories into 1-ns-long windows. Each nanosecond window is a time series of 200 locations of alpha carbon for each residue in MMP1. Correlations between every pair of residues are calculated for each window and normalized by dividing the correlation at lag number (time step) 0 (see methods and supplementary information about calculations of correlations). The plots (**Figures 3A-C**) show the matrix of mean correlation values over the entire simulated trajectory at lag (time step) number 1, normalized between 0 and 1. There are correlations in MMP1 dynamics, suggesting strong allosteric communications in MMP1.

**Figure 3.**
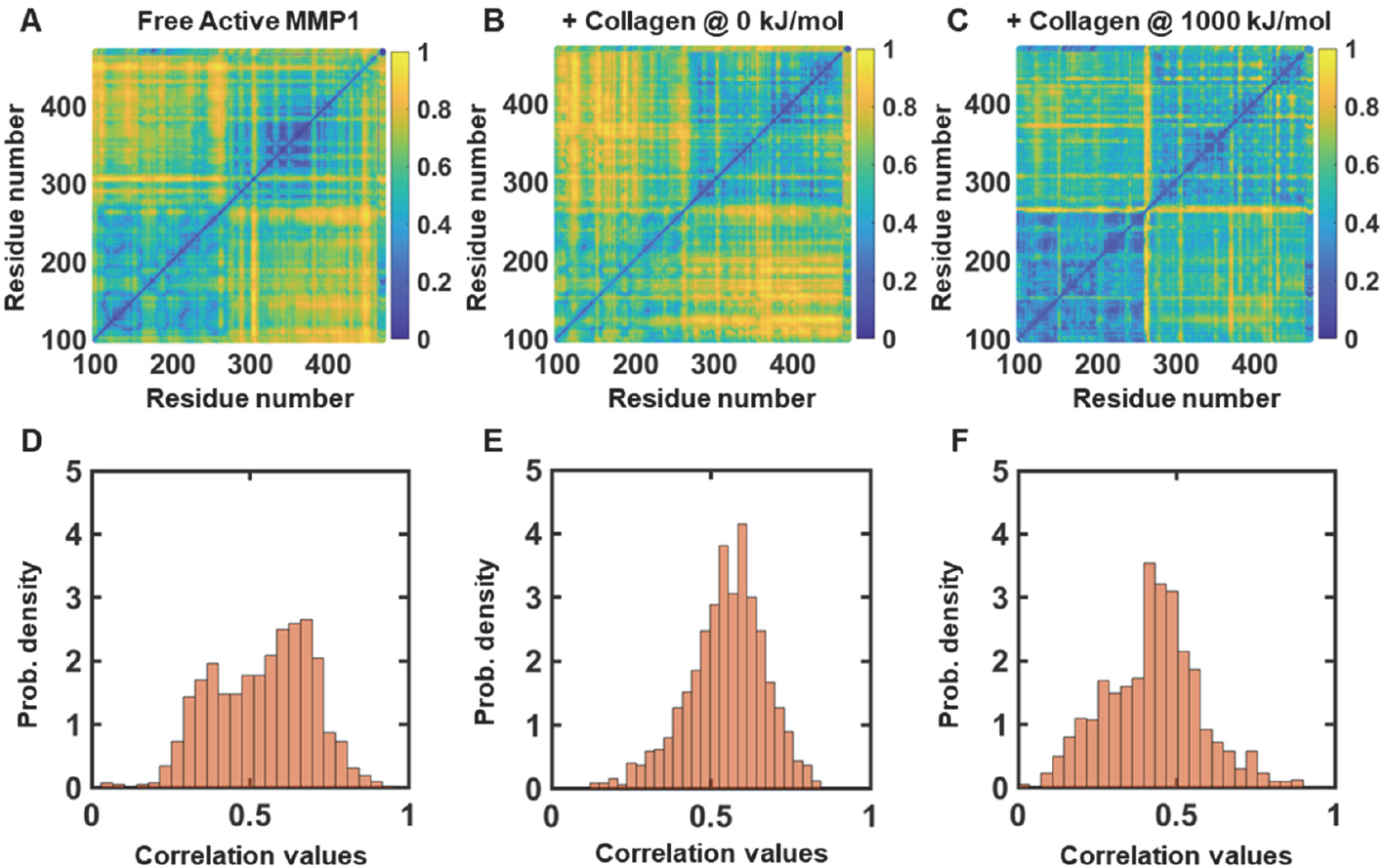
Correlation between each pair of residues in active MMP1. Pairwise correlation maps for (**A**) free MMP1, (**B**) collagen-bound MMP1 with 0 kJ/mol restraint on collagen, and (**C**) collagen-bound MMP1 with 1000 kJ/mol restraint on collagen. (**D-F**) Histograms of correlations in (A), (B), and (C).

### Allosteric residues in the hemopexin domain strongly correlate with the catalytic motif residues

Next, we plotted histograms of correlation values (**Figures 3D-F**) and selected the threshold correlation value determined by the mean ± two times the standard deviation for each model. We found all the residues whose normalized correlations to the 11 residues (HELGHSLGLSH) of the catalytic motif are greater than the threshold. We compared those residues between free MMP1 and collagen-bound MMP1 at 0 kJ/mol and 1000 kJ/mol.

We identified exclusive residues for Free MMP1 (30 residues: ARG291, GLY292, GLU293, THR305, ASN306, PRO307, PRO310, GLU311, VAL312, GLU313, GLY387, HIS417, GLY421, HIS424, LYS432, PHE435, LYS446, PHE447, ASP448, PRO449, LYS450, THR451, LYS452, ARG453, ILE454, LEU455, THR456, LEU457, ASN465, and CYS466), collagen-bound MMP1 at 0 kJ/mol restraint on collagen (30 residues: ASN346, LYS347, LEU357, HIS358, GLY359, TYR360, PRO361, LYS362, ASP363, ILE364, TYR365, SER366, SER367, PHE370, PRO371, HIS376, GLU384, LYS404, and ARG405), and collagen-bound MMP1 at 1000 kJ/mol restraint on collagen (8 residues: GLU154, SER263, GLN264, ASN265, PRO266, VAL267, GLN354, and GLY423).

Next, we investigated whether collagen-specific allosteric residues identified above have any potential functional relevance by sequence alignment. The hemopexin domain is known to regulate MMP1 catalytic activity (60). The hemopexin domain contributes to substrate and ligand specificity and activation and inhibition of various MMPs (61). The catalytic domain is similar across MMPs (11), but the hemopexin domain sequence varies (11, 62). This suggests that the evolutionary engineering of MMP activity occurred mainly via mutations in the hemopexin domain. We aligned the hemopexin sequences between ASP279 and CYS469 of MMP1, MMP14, and MMP9 from UniProt using ClustalW2. MMP1 and MMP14 can cleave type-1 triple-helical collagen, but MMP9 cannot. We found 32 amino acids between ASP279 and CYS469 that are the same in MMP1 and MMP14 but differ in MMP9. These mutations are D279N, R291G, G292N, E293Q, F320K, A331V, D336L, K344S, K347Q, G359P, I377V, D378T, A379G, T389M, Y390L, F391L, N395R, K396R, Y397L, E402V, R405Q, Y411S, P412A, K413S, I415V, I422V, M431Q, F435K, G441D, K446R, F447V, and W463I. Comparing the allosteric residues of collagen-bound MMP1 with the MMP9 mutations, we found R405 in MMP1 as the suitable candidate for functionally relevant allosteric residue that strongly correlates with the active site residues.

### Identified collagen-specific allosteric residue, R405, has allosteric paths connecting to the active site

We identified potential allosteric paths between R405 and the active site (**Figure 4**) using Ohm (63), a computationally efficient network-based method for protein allostery analysis. This tool employs percolation theory throughout the protein structure and is reliable for identifying and characterizing allosteric communication networks within proteins. Ohm uses a static tertiary structure as input and builds on the premise that protein structure is coupled to protein dynamics, and the allostery manifests this coupling. Further to this claim, the authors of Ohm propose that allosteric phenomena are a general property of heterogeneous media, and allostery propagates through regions of higher protein density.

**Figure 4.**
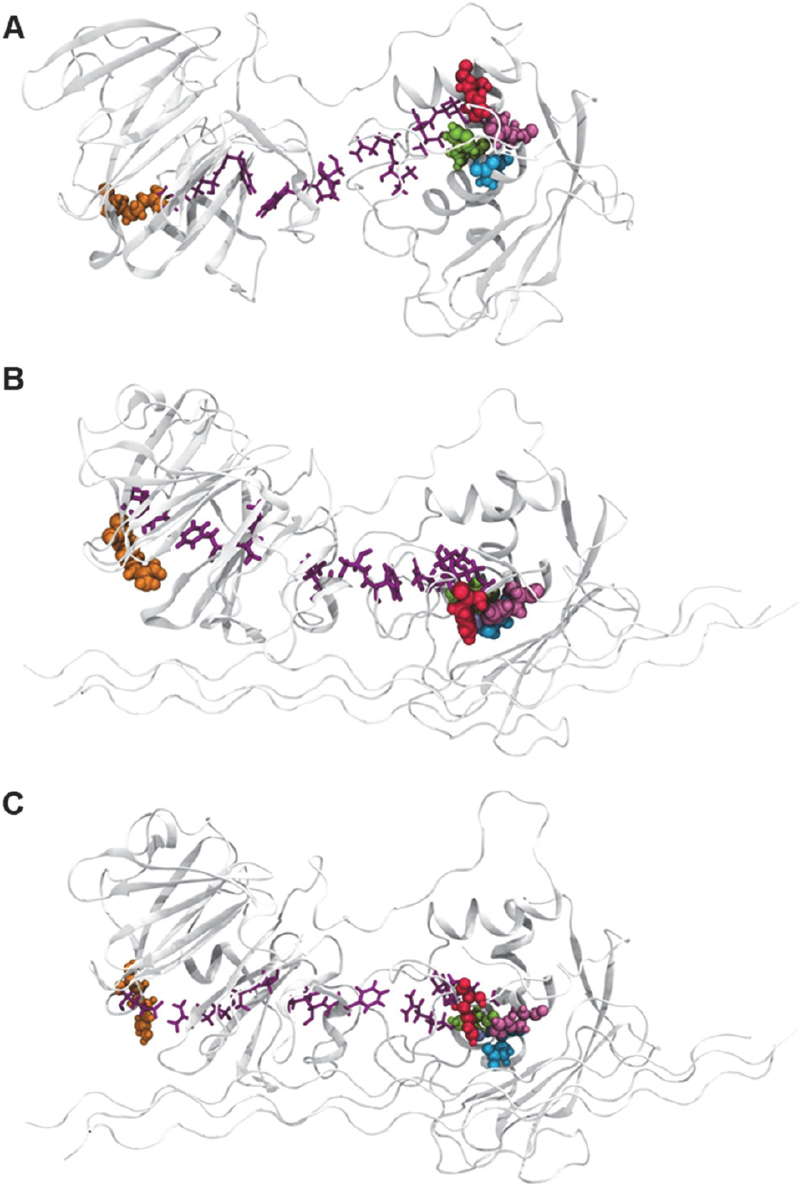
Potential allosteric paths from the allosteric residue R405 to the active site. The top-ranked allosteric pathway in MMP1 active (**A**) without collagen, (**B**) with unrestrained collagen, and (**C**) with 1000 kJ/mol position restraints on collagen. R405 is in orange spheres, the potential allosteric pathway is in purple sticks, and the active site is depicted in spheres and colored red, green, mauve, and cyan for H228, H218, H222, and E219, respectively. For the flowchart, see supplementary **Figure S3**.

We began by extracting representative frames from 1 μs MD simulations of active (E219) free MMP1, MMP1 with unrestrained collagen, and MMP1 with 1000 kJ/mol position restraints on the collagen alpha-carbon atoms. Simulations were performed following the energy minimization and relaxation steps outlined in the methods. We discarded the first five nanoseconds (for protein relaxation) from the production trajectory, and the remaining frames were subjected to cluster analysis using the Gromos method, with an RMSD cluster requirement of 0.1 nm on MMP1 heavy atoms. Representative frames were taken from the center of each cluster. These steps are illustrated in **Figure S3**. Water and counter ions were removed, and the final structures were submitted to the Ohm online service, totaling 30 submissions.

We defined the active site as H228, H218, H222, and E219 and the allosteric residue as R405. The residue distance cut-off was set to 0.4 nm, and the number of perturbation propagations to 20,000. Ohm predicted several potential allosteric pathways for each structure. **Figure 4** depicts the allosteric pathway with the largest allosteric coupling intensity (ACI) value for Free MMP1 (**Figure 4A**), MMP1 with unrestrained collagen (**Figure 4B**), and MMP1 with position restraints on collagen (**Figure 3C**). Allosteric paths from R405 to the active site differ for free and collagen-bound MMP1. Starting from R405, the top scoring allosteric path for free MMP1 is R405 → R341 → F342 → W321 → P322 → S318 → Q247 → V246 → L235 → M236 → the active site. For collagen-bound MMP1 with collagen restraint at 0 kJ/mol, the top scoring allosteric path is R405 → Y400 → T389 → Y332 → A331 → A330 → I317 → S243 → F242 → R214 → L235 → M236 → the active site. For collagen-bound MMP1 with collagen restraint at 1000 kJ/mol, the top scoring allosteric path is R405 → E402 → R341 → A331 → A330 → I317 → F316 → R214 → L235 → M236 → the active site. These results suggest that collagen changes the allosteric path in MMP1, and the allosteric paths depend on the collagen strain.

When we superimposed the top ten allosteric paths (**Figures S4-S6**), all pathways avoided the linker region and passed through the residues between the hemopexin and catalytic domain. Experimentally, a mixture of two MMP1 domains purified separately without the linker region can degrade collagen (5). Our previous paper suggested that collagen mediates the allostery between two domains (6). **Figure 4** shows that we need to discard collagen-mediated allostery without further proof and conclude that two domains can be allosterically connected via non-bonded interactions.

We noted differences when we superimposed the top-scoring pathways from each of the ten representative frames per model. Free MMP1 reveals a loose grouping of pathways, indicative of a greater degree of conformational freedom in the absence of collagen. Interestingly, we found unrestrained collagen models had a tight grouping of top-scoring allosteric pathways compared to when position restraints were introduced to the collagen substrate. In other words, we have shown that R405 is connected allosterically to the active site of MMP1, and collagen strain changes MMP1 allostery.

In light of collagen-mediated changes in dynamics, allosteric communications, and allosteric paths shown in **Figures 2-4**, we gained insights into the nature of allostery in MMP1-collagen interactions. MMP1 binds collagen strongly (64), and as such, we are likely in the strong perturbation regime when conformations (normal modes) of free MMP1 do not represent the conformations of the collagen-bound MMP1. Instead, we need to consider the whole system to identify conformational distributions. Indeed, the distributions of interdomain distance in **Figure 2** suggest that collagen does not merely select conformations of free MMP1; instead, collagen-bound MMP1 has an entirely different distribution of interdomain distance.

### Collagen binding leads to a significant change in the configuration of the MMP1 catalytic motif

There are three possible mechanisms of MMP catalysis at the active site. Browner et al. (65) suggested that the catalysis happens due to the conserved glutamic acid and the Zn^2+^ ion. Kester et al. suggested that catalysis happens due to interactions between the Zn^2+(66)^ ion and a water molecule, later proved unlikely by Manzetti et al. (67), who indicated that a histidine from the HEXXHXXGXXH-motif (HELGHSLGLSH for MMP1) allows the Zn^2+^ ion to assume a quasi-penta coordinated state. Five oxygen atoms (two from the catalytic glutamic acid, one from the substrate’s carbonyl oxygen, and two from histidine residues) can coordinate with the penta state of the Zn^2+^ ion, polarize the glutamic acid’s oxygen, proximate the scissile bond, and induce the scissile bond to act as the reversible electron donor. The resulting oxyanion transition state can facilitate a water molecule to act on the scissile bond to complete the catalysis. We studied how the substrate changes the catalytic motif configurations to gain more insights into the mechanism at the active site of MMP1.

Using 100 ns trajectories of the three models in their active form, we considered the catalytic residue E219 as the origin and plotted the center of mass for the catalytic motif residues. The plot-symbol size of the locations is proportional to the standard deviation of the pairwise distance (**Figure 5**). The configuration at the catalytic site changes considerably as free MMP1 binds collagen, and the fluctuations (plot-symbol sizes) reduce upon collagen binding. We recognize that the coordination of the catalytic zinc cannot change when using a model-bonded metal and classical force field implementation. While the catalytic zinc retains its penta-coordinated state, the interdomain distance was used to measure activity. We recognize this limitation, and we are exploring quantum-mechanical molecular models and artificial neural network-based all-atom models to increase the accuracy of our work. Nevertheless, **Figure 5** shows that the catalytic motif opens up more upon collagen binding than free MMP1, which is consistent with the fact that MMP1 must open up before the catalytic site gets closer to the collagen cleavage site (55, 68).

**Figure 5.**
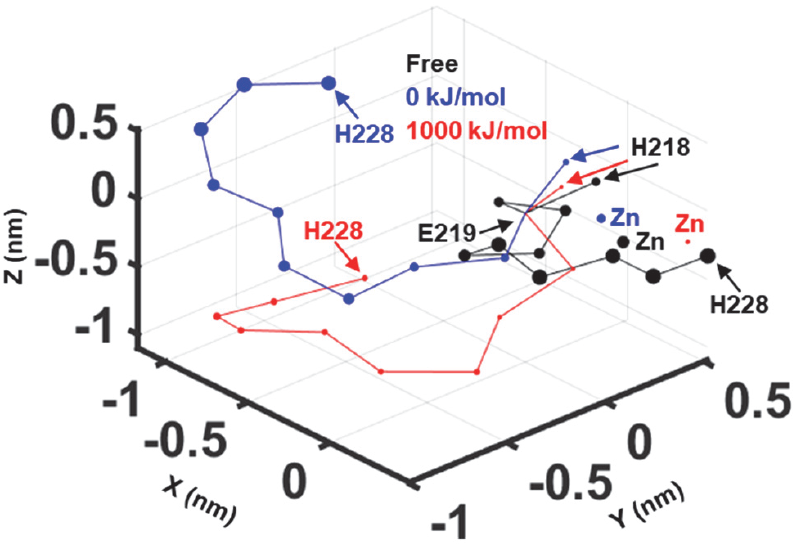
Collagen binding changes the catalytic motif configuration of MMP1. Three-dimensional configurations of the catalytic motif for free and collagen-bound MMP1 taken from 100 ns simulations. Residue positions are averages over the whole simulation length, with each respective configuration’s GLU219 placed at the origin. Symbol sizes are proportional to the standard deviation of the distance between the origin (E219) and individual catalytic motif residues. The data was obtained from the same simulations in **Figures 2** and **3**.

### Virtual screening of small molecules at the proposed allosteric site

One of our primary objectives was to conduct a virtual screen of single small molecules over the proposed allosteric site to understand their differences in binding affinities and how those binding affinities change when MMP1 is bound to the collagen substrate. Our meticulous process involved cataloging small-molecule binding affinity scores at the proposed R405 allosteric binding site of MMP1 in the presence and absence of the collagen substrate and how a change to the restraint on collagen alpha carbons affects the binding affinity. We then carefully selected suitable candidates for further analysis, adjusting constraints on collagen to understand the effect substrates have on small-molecule binding at allosteric sites.

A small bounding search box was centered at the center-of-mass of R405, located on the surface of the hemopexin domain. Polar hydrogens and Gastiger charges were applied. “AutoDock-ready” PDBQT small molecule structures for immediate turnkey screening were taken from ZINC 22 (69). We randomly selected 0.05% from each tranche that makes up the Goldilocks selection (approximately 140,000 small molecules). Suitable small molecule ligands for high-throughput screening are characterized by a molecular weight between 250-500 Daltons, a LogP between 1 and 3, a moderate number of rotatable bonds (usually less than 8), a polar surface area of less than 140 angstroms, and overall, adhere to Lipinski’s Rule of Five (70).

We performed two virtual screens using AutoDock Vina with the preferred default settings (as per the “turnkey” implementation) and those documented in the methods. The first virtual screen was performed on the MMP1 crystal structure (4AUO) with collagen. The second virtual screen was performed on MM1 without collagen. A 100 ns NPT simulation of active MMP1 without collagen was performed, and we retrieved the median frame from the largest cluster (using the Gromos method) as the representative frame for virtual screening.

The top score from each small molecule was cataloged (**Figure 6A**), and the 3D coordinates of the pose were recorded. Our calculations suggested that small molecules could bind to MMP1 without collagen (the average was approximately −25 kJ/mol) more so than in the presence of collagen (the average was approximately −22 kJ/mol). The top-scoring pose per small molecule returned a negative binding affinity.

**Figure 6.**
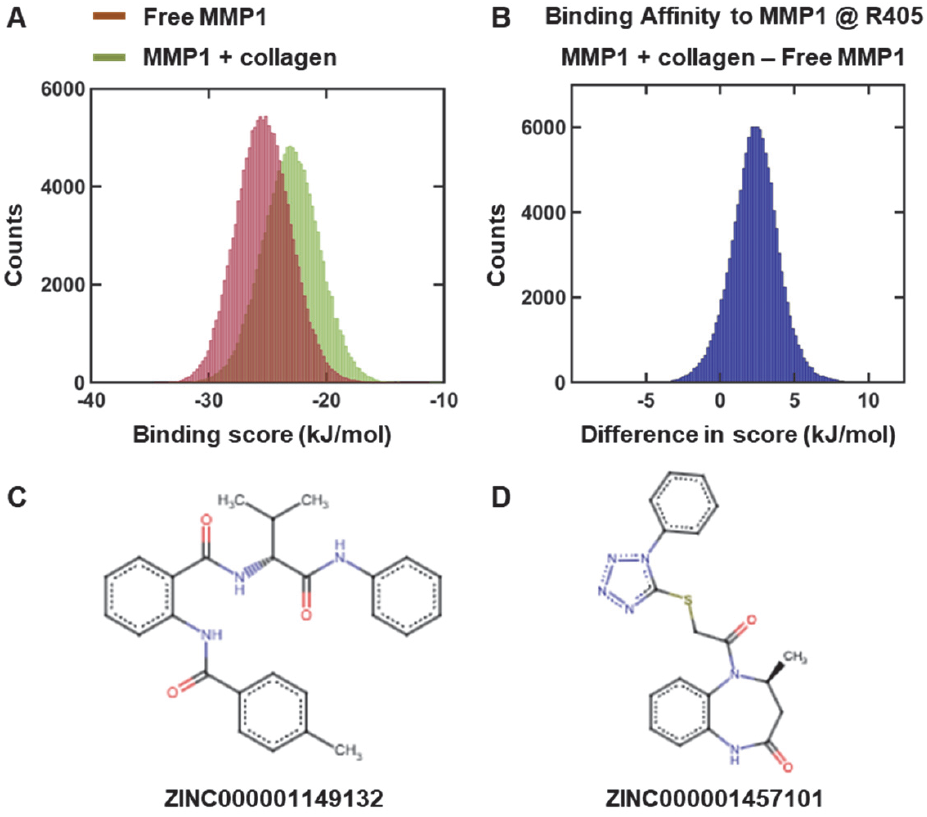
The binding affinity scores from a small molecule virtual screen against MMP1. (**A**) Small molecule binding affinity scores against Free MMP1 and MMP1 with collagen. (**B**) Binding affinity score difference for 140,000 small molecules using AutoDock Vina between free and collagen-bound MMP1. We selected two molecules: (**C**) ZINC000001149132 (ZINC11) and (**D**) ZINC000001457101 (ZINC14).

Having performed the two virtual screens, we calculated the difference in binding affinity score between the top pose from each small molecule when screened over MMP1 with and without collagen (**Figure 6B**). Most binding affinity scores became positive, indicating Free MMP1’s lower binding affinity scores. This underscores the importance of our research. We did not aim to find small molecules with the most significant binding affinity to one target structure but to look for the small molecule with the most significant positive difference and one with the significant negative difference.

We took the compound with the most positive difference **(Figure 6B)** in binding affinity (ZINC000001457101) at 12.388 kJ/mol and the largest negative difference in binding affinity (ZINC000001149132) at −5.149 kJ/mol. When retrieved from the virtual screen in the presence of collagen, ZINC14 and ZINC11 returned binding affinities of −21.781 kJ/mol and −29.087 kJ/mol, respectively. When screened without collagen, the corresponding binding affinities switched, with −34.170 kJ/mol and 23.938 kJ/mol, respectively. Although ZINC11 and ZINC14 bind to free and collagen-bound MMP1, their binding scores differ. ZINC11 binds stronger to free MMP1, whereas ZINC14 binds stronger to collagen-bound MMP1.

We recognize the limitations of these small-docking calculations; AutoDock Vina estimates binding affinity using a combination of terms that approximate intermolecular interactions. However, the scores do not account for long-range electrostatics, solvation dynamics, or entropy (71). Therefore, we took the ZINC11 and ZINC14 poses and rescored them after free energy calculations using MD simulations.

### Small molecule binding affinities to MMP1 depend on the substrate

To assess the impact the collagen substrate might have on the underlying allosteric dynamics in MMP1, we compared the free energies of binding at the allosteric site of two small molecules from our AutoDock Vina screens (**Figures 7A** and **7D**) against the three MMP1 active models: Free MMP1, MMP1 with unrestrained collagen, and MMP1 with 1000 kJ/mol position restrained applied to the collagen alpha-carbon atoms.

**Figure 7.**
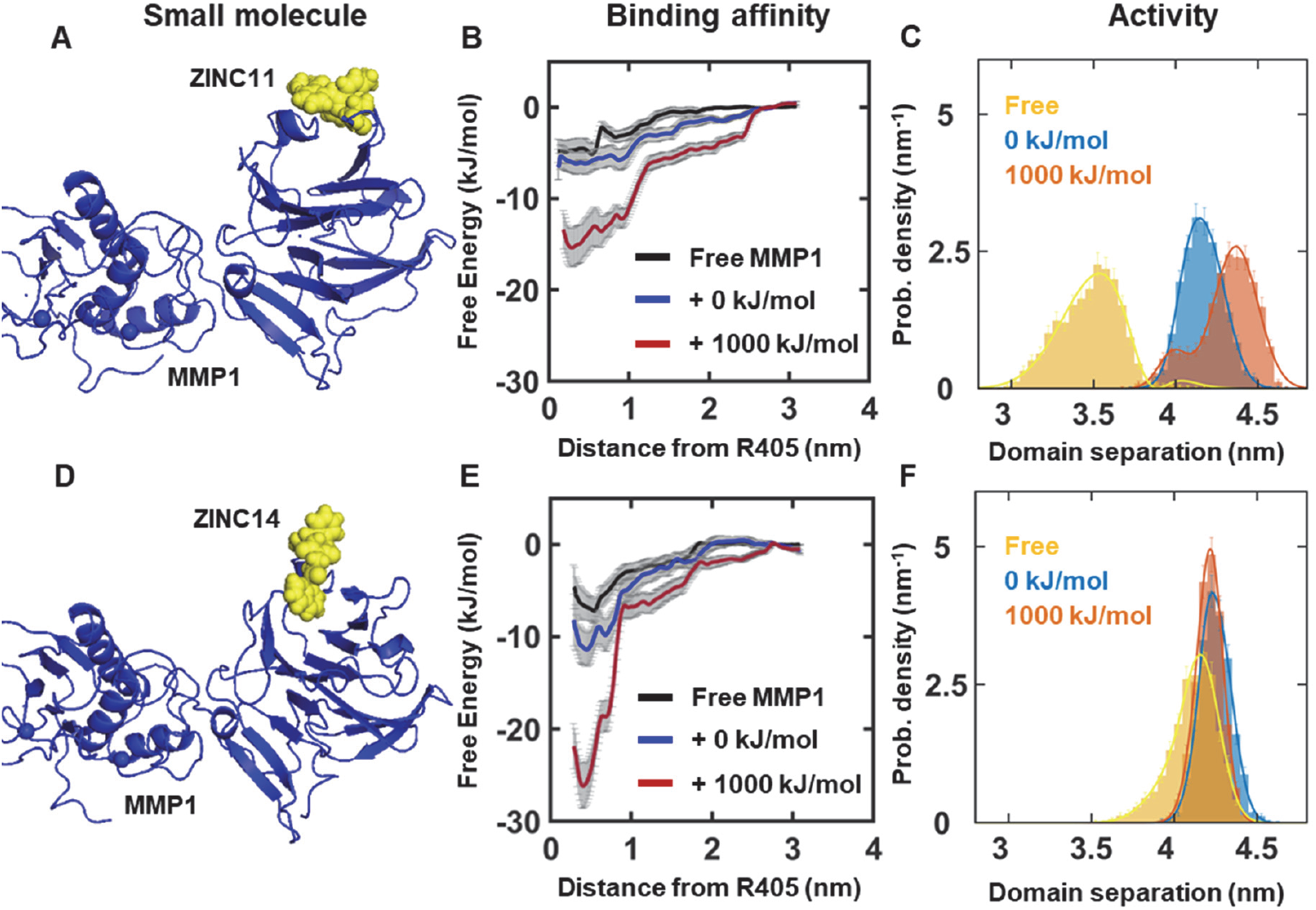
Small molecule binding affinities to MMP1 depend on the collagen flexibility. (**A**) A representative pose of ZINC11 bound to free MMP1. (**B**) Free energy of ZINC11 binding. (**C**) Interdomain distance when MMP1 is bound to ZINC11. (**D**) A representative pose of ZINC14 bound to collagen-bound MMP1. (**E**) Free energy of ZINC14 binding presented. (**F**) Interdomain distance when MMP1 is bound to ZINC14. The data for **(C)** and **(F)** was obtained from 100 ns simulations with 20 ps time resolution. The starting pose before the relaxation of these simulations was the best binding pose found by AutoDock Vina.

**Figure 8.**
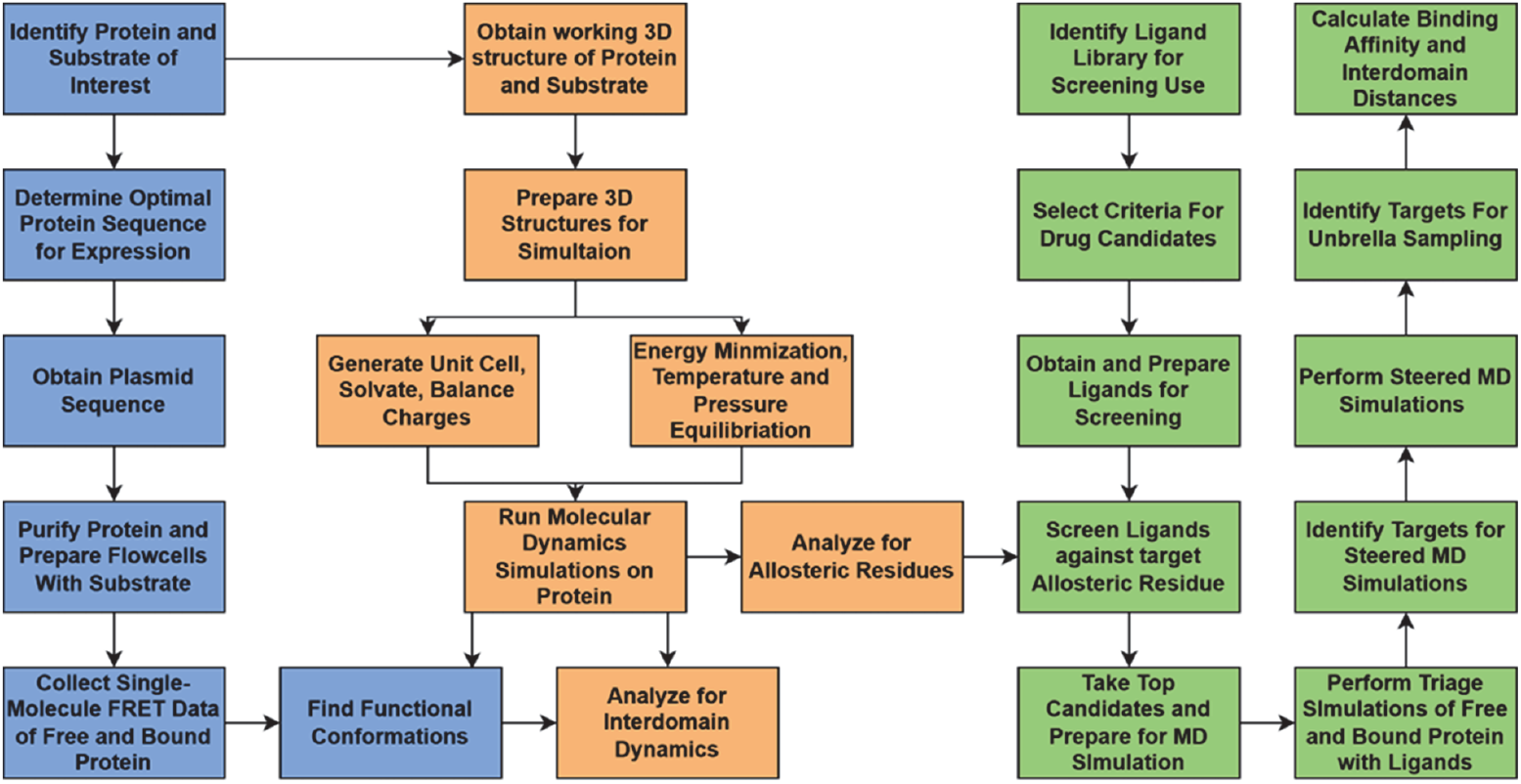
A framework for predicting small molecules to achieve substrate-specific control of enzyme activity. First (in blue), we perform experiments to identify functional conformation. Second (in orange), we perform simulations, validate with experiments, and identify substrate-specific allosteric residues. Third (in green), we select small molecules from the ZINC database, screen against the allosteric residues, perform MD simulations, calculate the binding affinity of the small molecules to the target protein, and calculate the interdomain distance to correlate with activity. We confirm that the selected small molecules have stronger binding affinity than other locations on the surface of the targeted protein. Finally, we experimentally validate the effects of identified molecules.

We performed a steered MD simulation using the NPT ensemble to pull the small molecule off the protein’s surface into the bulk solvent with a pull force constant of 1000 kJ/mol/nm^2^ and a pull rate of 0.3 nm/ps. We aimed to achieve a clearage of at least 3 nm from the original docked pose, a crucial step to ensuring the ligand reaches a state where it no longer interacts with the protein, effectively representing an unbound state, clearing non-bonded interactions and ordered water-mediated effects. The trajectory became a linear reaction coordinate. Although umbrella sampling along a linear reaction coordinate is unlikely to reveal a realistic binding pathway, umbrella sampling estimates the difference in free energy between the bound and unbound states without being path-dependent.

Independent frames were taken every 0.1 nm along the reaction coordinate. Each frame was subjected to a steepest descent energy minimization process, relaxed as discussed in the methods, and simulated for 40 ns with a pull coordinate rate of 0.0 nm/ps. This was performed three times to ensure sufficient coverage along the reaction coordinate (**Figure S7**) and maximize independent conformational space sampling. We recalculated each PMF calculation for convergence while increasing the sample size (**Figure S8**). We shifted the PMF plots so the unbound state was 0 kJ/ mol.

The free energies for both small molecules against the three models were negative **(Figures 7B** and **7E)**, suggesting a strong propensity to bind at the site approximal to the allosteric residue. The Free MMP1 model yielded the weakest non-bonded association for both small molecules, with ZINC11 (PMF minimum of −5.02 kJ/mol) 2.13 kJ/mol weaker than ZINC14 (PMF minimum of −7.15 kJ/mol), however, they were within error. These results were closely followed with unrestrained collagen. The free energy of binding for ZINC11, −6.17 kJ/mol, was within error of the Free form, while ZINC14 yields a significant decrease to −11.2 kJ/mol. When 1000 kJ/mol are applied to the collagen, the PMFs for both small molecules show a substantial reduction in free energy, −15.4 kJ/mol for ZINC11 and as much as −26.1 kJ/mol for ZINC14. Although screening software such as AutoDock Vina (72) currently provides a valuable tool to downselect small molecules from an extensive database, they do not consider dynamics and the effects of target protein’s substrates. They should be used with additional methods for accurately identifying potential lead molecules.

## Conclusion

In conclusion, we have used validated MD simulations to gain insights into collagen-mediated changes in MMP1 dynamics, allostery, and ligand binding. We have found that strain in collagen affects MMP1 dynamics, consistent with previously published experiments on strain-dependent activity of MMP1 (42) and age-dependent changes in collagen strain in humans (53). Simulated dynamics match with experimental dynamics only when we impose a restraint on collagen alpha carbons. Validated simulations clearly show that the interdomain separation of MMP1 increases with collagen strain, which we showed to be functionally relevant in our previous paper (6). Researchers support (73) and refute (74) the essential role of conformational dynamics in activity. Our results support the functional role of MMP1 dynamics and are consistent with the experimental observation that both domains are necessary to degrade triple-helical collagen (5).

Simulations of MMP1-collagen interactions revealed insights into allostery in MMP1. Residues in the hemopexin domain have strong correlations with those in the catalytic domain, and we found collagen-specific allosteric residues or “fingerprints” in the hemopexin domain. A sequence alignment showed one of the identified allosteric residues, R405, is mutated to glutamine in MMP9, which cannot degrade type-1 triple helical collagen. We identified the allosteric paths from R405 to the active site of MMP1, which showed no paths through the linker domain of MMP1. Instead, there are non-bonded interaction bridges between the allosteric paths in the two domains.

We screened small molecules against the collagen-specific allosteric residue, R405, using AutoDock Vina (72) for free and collagen-bound MMP1. We identified two molecules with the highest binding scores; one prefers free MMP1, and the other prefers collagen-bound MMP1. However, when we calculated the binding affinities using MD simulations, both small molecules preferred collagen-bound MMP1 with 1000 kJ/mol restraint on collagen alpha carbons. These results suggest we should consider the substrate while screening molecules against a protein. Although currently available screening packages, such as AutoDock Vina (72), provide tools to down-select small molecules from an extensive database, they do not consider the dynamics and effects of the target protein’s substrates. As such, these screening tools must be supplemented with additional methods to identify potential lead molecules accurately.

To summarize, we have gained fundamental insights into the role of substrate in a protein’s dynamics, allostery, and ligand binding. We have also developed a framework for screening small molecules against substrate-specific allosteric residues in MMP1 by combining single-molecule experiments, all-atom MD simulations, and virtual screening.

## Materials and methods

For this study, we have built several models based on the active (E219) and catalytically inactive point mutant (E219Q) of human MMP1. We started with PDB ID 4AUO (41) and constructed free and collagen-bound models of human MMP1 using a collagen-like triple-helical substrate. We used AutoDock Vina (72) to build two sets of protein-drug complexes, specifically ZINC000001149132 (ZINC11) and ZINC000001457101 (ZINC14), with the active form of free and collagen-bound MMP1. Below, we describe how we built each model, followed by the MD parameters, the virtual screen, and ligand binding calculations. Finally, we will explain how we calculated the free energy of ligand binding. MD simulation length and equilibration steps are described throughout the results.

### MMP1 simulation model construction

We began model construction using the PDB-redo corrected form of inactive human MMP1 bound to a collagen-like triple-helical substructure (PDB ID 4AUO) (41). MMP1 atoms were encoded using the AmberSB99ILDN forcefield, and GAFF2 topology files were built from ligand PDBQT files using AmberTools Antechamber. Acetyl group and methylamine group residues were used to terminate the MMP1 molecule, and when present, collagen was modeled as a periodic molecule across the periodic boundaries. The periodic-collagen molecule was relaxed using an unrestrained 100 ns simulation using the NPT ensemble to adjust the unit cell and collagen atoms to the periodic bond introduced to the collagen backbone.

We built the active form by substituting A219 (A200 in the PDB) for E219. We used the force field parameters from our previous published work for the two zinc and four calcium ions (6). In brief, the parameters were calculated using Amber’s Metal Center Parameter Builder (MCPB.py) (75), and electrostatics were calculated using RESP (restrained electrostatic potential) charges. All models were solvated using tip3p water, and the total system charge was neutralized using counterions.

The two active ligand-MMP1 complexes, with collagen substrate and free, were prepared using the lowest binding score form of the corresponding ligands, as described further in the methods. We translated and rotated the ligand structure from the coordinate space of AutoDock Vina to a prepared MMP1 protein simulation model using MDAnalysis (76, 77) and MDLovofit (77). Regions of low mobility (approximately no more than 1 angstrom) were identified and used as an anchor to calculate a transformation matrix from the corresponding docking receptor protein onto the MD model. This step was required to preserve protein-ligand contacts. The simulation models had been subjected to a series of pre-simulation adjustments, including reorientation to account for the periodic triple helical collagen molecule, solvation, and energy minimization calculations.

### MMP1 simulation parameters

We used Gromacs 2023.4 (78) to integrate the atomic step using the Velocity Verlet algorithm with a time step of 0.02 ps and recorded coordinates and velocities every 5 ps. Bond constraints were applied to hydrogen atoms using the LINCS schema, with the LINCS iteration set to one and with an order of four. We controlled and updated the cut-off of the neighbor search list every ten steps using the Verlet scheme. Periodic boundary conditions were applied in all directions.

We controlled long-range electrostatic interactions using the fast, smooth Particle-Mesh Ewald scheme, with an interpolation order of 4 and a Fourier spacing of 0.16 nm. Short-range energy interactions were controlled using cut-off Van der Waals interactions and short-range Coulomb interactions, set to 1.0 nm and modified with a potential shift.

The temperature was set to 295 K, and the temperature of all solvent and non-solvent atoms was controlled using the V-rescale thermostat with a temperature coupling of 0.1 ps. The pressure coupling was set to 1 atmosphere and was controlled using the Berendsen, followed by the C-rescale isotropic pressure coupling with a time constant of 2 ps. The compressibility was set to 4.5e-5 bar^−1^.

### Distance-based Analyses

We performed several calculations that required Euclidean distance measurements. The interdomain distances between the center-of-mass of S142 in the hemopexin domain and the center-of-mass of S366 in the catalytic domain were calculated using the Gromacs gmx pairdist function. We picked these residues to reflect our previously published work in which they were mutated to attach a dye label for single-molecule Förster resonance energy transfer. Periodic conditions were resolved to avoid distance artifacts, and distance distributions were compiled in MATLAB using *in-house* scripts.

We used the RMSF of the collagen alpha-carbon to measure the flexibility of individual MMP1 residues. To achieve this, we used MDLovofit (77) to identify the protein fraction in which the RMSD of the alpha carbons was less than 1 angstrom (the least mobile residues). The MMP1 protein was iteratively repositioned over the region with the lowest RMSD, and the RMSF of the alpha-carbons was calculated. This ensured we could control for domains with sizeable spatial displacement.

### Analyses of experimental and simulated dynamics

We quantified dynamics by conformational histograms and correlations, as we did for previous publications (6, 79). For histograms, we chose bin widths equal to or larger than 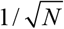, where *N* is the number of data points. We calculated each bin’s error as the bin count’s square root. To obtain the probability density function for conformations, we divided the bin counts and errors by the histogram area so that the area under the normalized histogram equals 1. We fitted a sum of two Gaussians to the area-normalized histograms to determine the two primary conformational states of MMP1 and their widths:

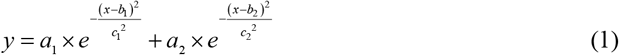

where a’s, b’s, and c’s are amplitudes, centers, and widths of the Gaussians. Two Gaussian centers, b1 and b2, are the two states, S1 and S2, of MMP1.

For correlations among conformations, we used the following equation:

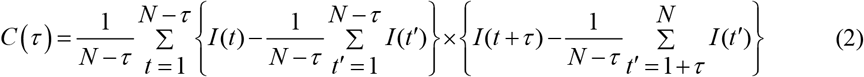

where *C* (*τ*) is the correlation at lag number *τ, N* is the number of points in a FRET time series, and *I* (*t*) is the FRET value at *t*. For autocorrelations, both factors in curly brackets were from the same time series. For cross-correlations, the two factors in curly brackets were from different time series representing coordinates of different residues. We normalized correlations by dividing correlation values at each lag by the correlation value at *τ* = 0. For an example of calculation and further explanation, see the supplementary information on “Calculation of correlations.”

### Allosteric Pathway Analyses

We used the Ohm online service to identify a potential allosteric pathway between the proposed allosteric binding site (proximal to R405) and the three histidine residues: H218, H222, and H228, and the glutamic acid, E219, coordinating with the zinc ion in the catalytic active site. We submitted frames isolated using the Gromos cluster analysis, as implemented in the Gromacs gmx cluster function. The radius cut-off to define cluster members was set to 1 Å. A representative frame from the ten largest clusters was submitted to the Ohm server. We set the distance cutoff of contacts to 4 angstroms, the alpha-value to 3 (default), and 20,000 rounds of perturbation propagation to identify allosteric pathways. Finally, we identified the top potential allosteric pathway by its measure of the “importance of allosteric pathway.”

### Virtual screen and ligand binding calculations

From the extensive pool of 270 million “Goldilocks” ligands (LogP up to 2 - 3, and a molecular weight of 300 - 375 Daltons) in the ZINC 22 database (69), we randomly selected 0.05 % from each chemical tranche, resulting in a total of 140,000 ligands.

We performed a virtual screen on two MMP1 protein targets. Firstly, we isolated an MMP1 free representative frame from non-restrained MD simulations of the Redo 4AUO crystal structure by generating structural clusters based on coordinates of the MMP1 alpha-carbons using the gromos algorithm (part of the Gromacs gmx cluster) across a 200 ns MD simulation trajectory with a 2-Å RMSD cut-off. A representative structure is a frame with the smallest average distance to other structures in the cluster with the greatest number of frames. For free MMP1, we used the Redo crystal structure of MMP1 with adjustments to several key residues to reflect our experimental conditions (mentioned above). The system was subjected to an energy minimization procedure using Gromacs.

We used the turn-key AutoDock Vina program (version 1.2.5) (72) to screen 140,000 ligand molecules. Each protein target was appropriately protonated with polar hydrogens, assigned Gastiger chargers, and AutoDock atom types using AutoDock Toolkit 1.5.7. The grid cube was centered on the carbon-alpha of R405 (R386 in the PDB), and grid spacing was kept at the default of 0.35 angstroms, set to 25 spaces across the three cube dimensions. When collagen was present, the substrate was beyond the grid cube.

Two lists of binding affinity scores were ordered by their ZINC ID. The difference in binding affinity score between MMP1 with collagen and free was calculated and reordered by the absolute difference in binding affinity score. The ten ligands with the largest absolute difference in binding score were rescored using atomistic MD free energy calculations.

### Free energy of ligand binding calculations

Before rescoring ligand binding using free energy calculations, we performed a ten-ns unrestrained MD simulation using the ten ligands with the greatest binding score between the collagen-bound and free forms. This step helps control solvation and entropic effects once the bound-ligand candidate forms are converted into MD simulations. The ligand-protein complex was relaxed following the steps described in the simulation details section. Simulations in which the ligand was shown to disassociate from the protein were discarded. The two model pairs (with collagen and free) with the largest absolute binding score difference and remaining in place during the simulation were subjected to a steered MD (SMD) simulation.

We used SMD to define a reaction coordinate for binding free energy as the Euclidian distance in 3D coordinate space from the center of geometry of the three nearest MMP1 alpha-carbons to the ligand and a point in bulk solvent. The direction was carefully chosen to ensure the ligand reached the bulk solvent while avoiding collision with the protein. Although a straight reaction coordinate is unlikely to represent the binding pathway, we can use it to calculate an approximate binding free energy. The reaction coordinate was defined using a direction-based umbrella harmonic potential between the proposed binding position (described above) and the ligand center of mass. A pull force constant of 1000 kJ/mol/nm^2^ with a pull rate of 0.3 nm/ps was applied to the direction pull coordinate.

Individual simulation windows were extracted from the SMD trajectory every 1 angstrom along the reaction coordinate. The pull coordinate rate was reduced to 0.0 nm ps^−1^, and each simulation window was energy minimized. This was followed by a 40 ns NPT simulation performed in triplicate (independent velocities), totaling 120 ns per simulation window. We recorded forces along the reaction coordinate every ten picoseconds. The first two nanoseconds were discarded from each simulation.

To construct potential mean force plots (PMF) following the reaction coordinate of ligand binding, we used the Gromacs implementation of the weighted histogram analysis method (WHAM) (80). Statistical uncertainty was calculated using a Bayesian bootstrap approach with 500 samples. We carefully inspected the distribution of histogram points along the reaction coordinate to ensure sufficient spatial sampling. We also checked PMF plots using progressive collections of points in 5 ns increments for convergence. These are available in the supporting information.

We calculated the difference between bound (ligand-protein complex) and unbound (ligand in bulk solvent) by adjusting the PMF such that the point of PMF convergence into bulk solvent was 0 kJ/mol. Therefore, the ligand binding free energy value is the difference between the point of bulk solvent convergence and the lowest point on the PMF curve.

## Acknowledgments

This work was supported by a grant to S.K.S. from the National Institutes of Health (GM145210).

## Author contributions

S.K.S. conceived and designed the overall project. A.N. and C.H. performed simulations. A.N., C.H., and S.K.S. analyzed the results. S.K.S., A.N., and C.H. wrote the manuscript. All authors edited the manuscript.

## Supplementary information

### Quality checks and durations of simulations were used in this paper

We followed the best practices in MD simulations (1). We ensured the quality of the simulations before analyzing the data and making scientific conclusions. First, we monitored each model’s potential energy, temperature, and pressure to ensure the system, thermostat, and pressure coupling looked stable. Then, we checked the Root-Mean-Squared Deviation (RMSD) to ensure the stabilization of simulations. We defined the input structure at t=0 as the reference structure and calculated the RMSD.

**Figure S1.**
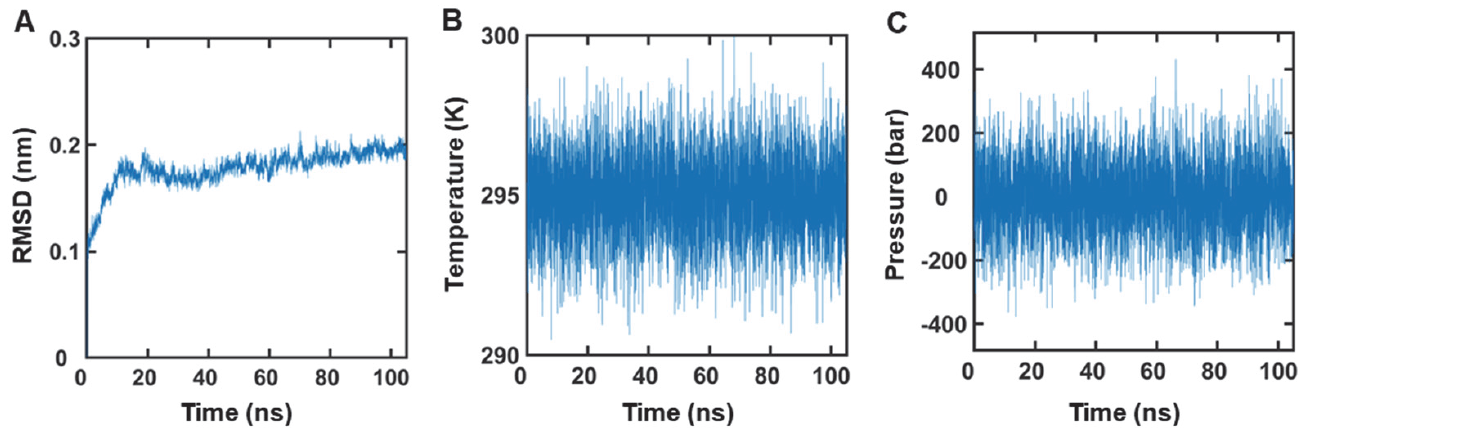
Quality checks of simulations. (**A**) RMSD, (**B**) temperature, and (**C**) pressure from simulations of collagen-bound MMP1 with collagen at 1000 kJ/mol position restraint on collagen alpha carbons.

For each model in **Figure 2** of the main text, we simulated 105-ns-long simulations and discarded the first 5 ns (**Figure S1**). For **Figures 2, 3**, and 5, we used the same six 105-ns-long simulations for a total simulation time of 630 ns. For **Figure 4**, we simulated three 1-μs-long trajectories for free MMP1, collagen-bound MMP1 with 0 kJ/mol restraint on collagen alpha carbons, and collagen-bound MMP1 with 1000 kJ restraint on collagen for a total simulation time of 3 μs. For **Figure 7**, we simulated three independent repeats of 40-ns-long simulations every 0.1 nm along the reaction coordinate for free MMP1, collagen-bound MMP1 with 0 kJ/mol restraint on collagen alpha carbons, and collagen-bound MMP1 with 1000 kJ restraint on collagen, for a total simulation time of 21.6 μs. The data presented in this paper include a total of 25.23 μs simulations. For free MMP1 in water, we had 70,874 atoms, and we could run simulations for 116.16 ns per day using the Arizona State University supercomputing facility. For collagen-bound MMP1 in water, we had 63,741 atoms, and we could run 312.49 ns per day.

**Figure S2.**
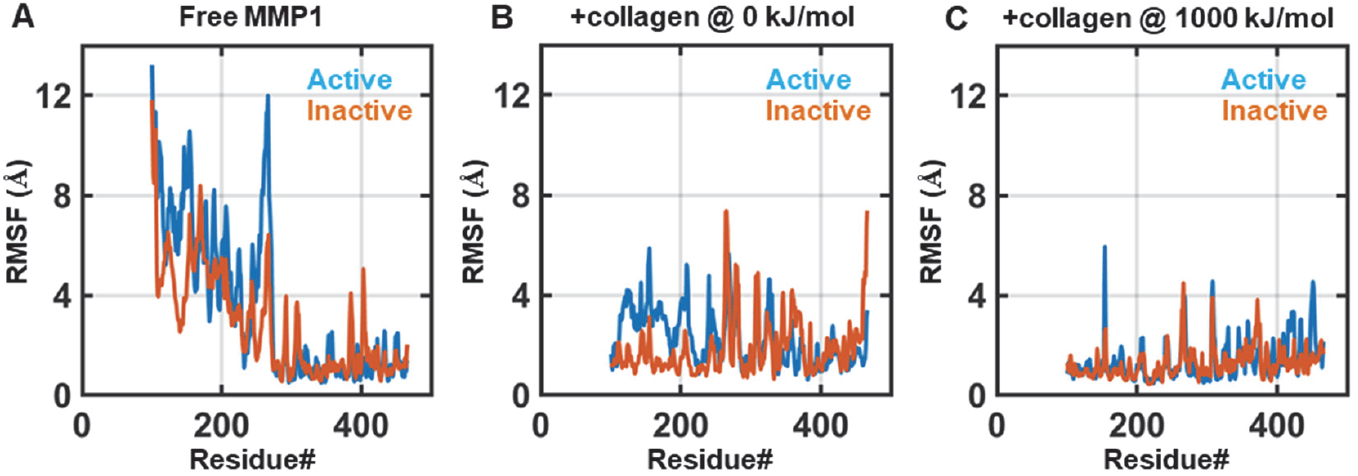
Fluctuations of individual MMP1 residues. RMSF of alpha carbon positions of residues for active (blue) and inactive (orange) for (**A**) free, (**B**) with collagen at 0 kJ/mol position restraint on collagen alpha carbons, and (**C**) with collagen at 1000 kJ/mol position restraint on collagen alpha carbons.

### Calculation of correlations

We used the following equation to calculate autocorrelations:

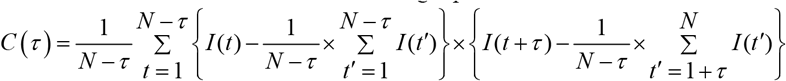

Where *C* (*τ*) is the correlation at lag number *τ, N* is the number of points in a time series, and *I* (*t*) is the value at *t*. We illustrate the process using a simple time series (*t,x*):

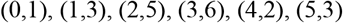

*C* (*τ*) at lag number*τ* = 0 :

In this time series, *N*=6 and the mean=(1+3+5+6+2+3)/6=20/6=3.3. After subtracting the mean to analyze only the fluctuations, we obtain a new time series:

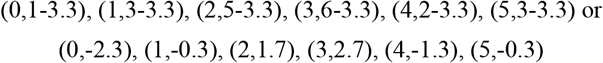

We do not need to shift the time series to calculate the correlation value for *τ* = 0 and can multiply element by element and calculate *C* (*τ*) at the lag number*τ* = 0 as below:

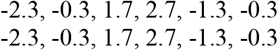

We multiply element by element, add, and average by *N* −*τ* =6−0 = 6 to get:

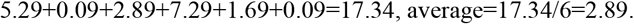

Therefore, *C* (*τ*) at lag number*τ* = 0 is 2.92.

*C* (*τ*) at lag number*τ* =1:

For *τ* =1, we need to shift the time series by 1 time step and multiply element by element:

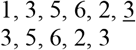

We disregard the last element, underlined 3, in the first row because it is unpaired. For the first row, the mean is (1+3+5+6+2)/5=17/5=3.4. For the second row, the mean is (3+5+6+2+3)/5=19/5=3.8. After subtracting the means, we get the following time series:

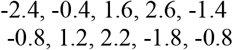

We multiply element by element, add, and average by *N* −*τ* =6−1=5to get:

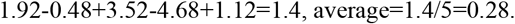

Therefore, *C* (*τ*) at lag number*τ* =1 is 0.28.

*C* (*τ*) at lag number*τ* = 2 :

For *τ* = 2, we need to shift the time series by 2 time steps and multiply element by element:

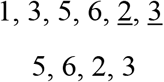

We disregard the last two elements, underlined 2 and 3, in the first row because they are unpaired. For the first row, the mean is (1+3+5+6)/4=15/4=3.75. For the second row, the mean is (5+6+2+3)/4=16/4=4.0. After subtracting the means, we get the following time series:

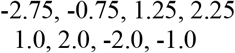

We multiply element by element, add, and average by *N* −*τ* =6−2 = 4to get:

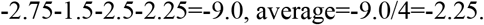

Therefore, *C* (*τ*) at lag number*τ* = 2 is −2.25. The autocorrelations at different lag numbers are:

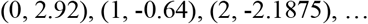

After normalization by dividing the value at *τ* = 0, we get:

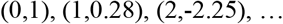

We have followed the procedure above to calculate normalized correlations.

**Figure S3.**
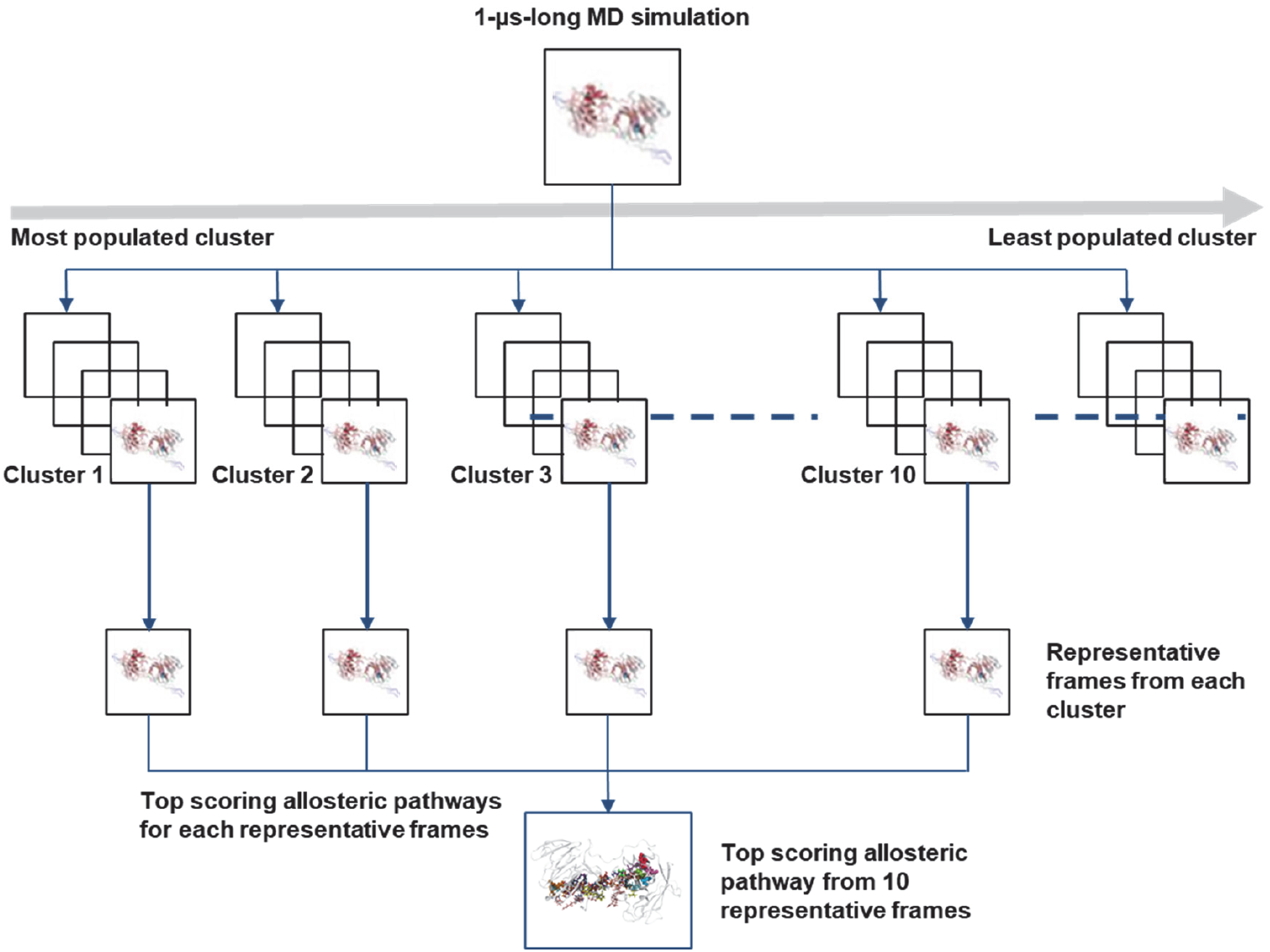
Workflow used to yield top-scoring allosteric pathways. We took representative frames from 1 μs MD simulations from the top ten clusters (ranked by cluster size). For each representative frame, the top-scoring allosteric pathway was retrieved from Ohm. The top-scoring pathway from the ten structures is presented in the main body of the manuscript, and the top-scoring pathway from each cluster is presented here in the SI.

**Figure S4.**
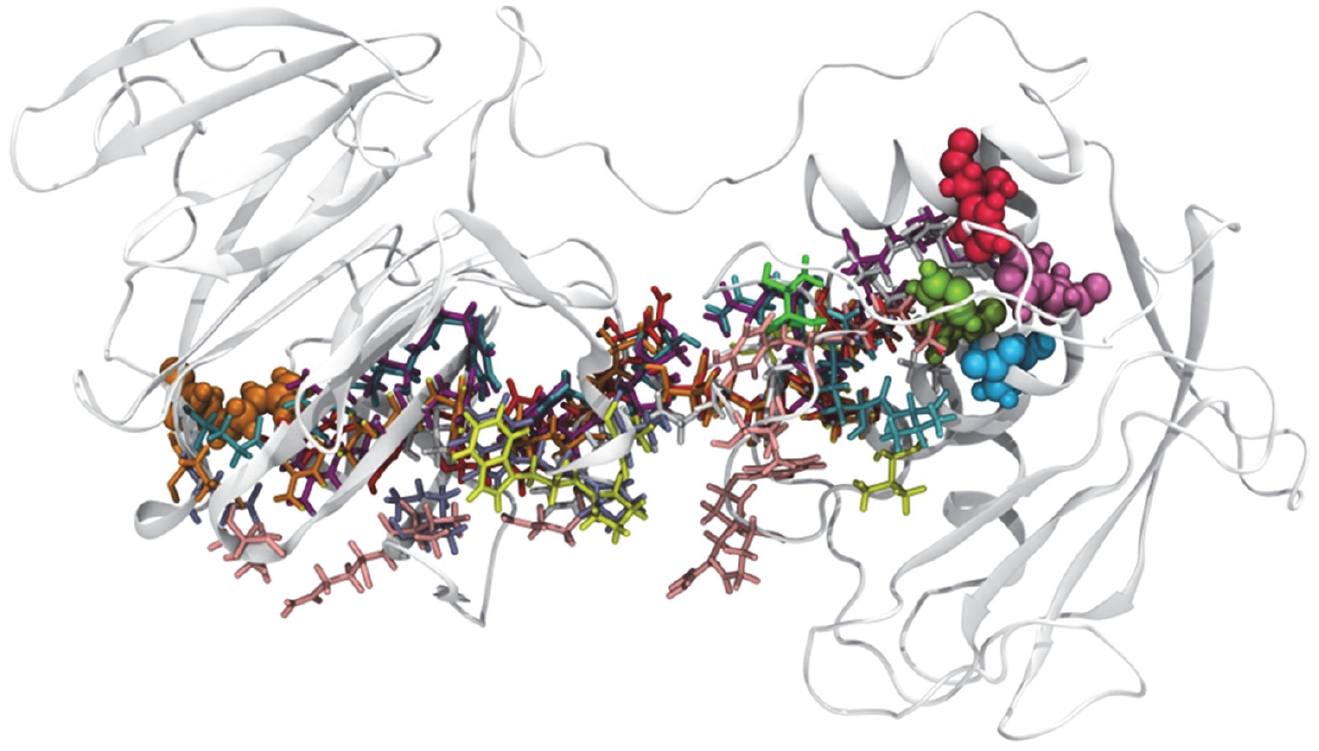
Pathways depicting the top allosteric coupling intensity values from free MMP1. Singular pathways are represented by color.

**Figure S5.**
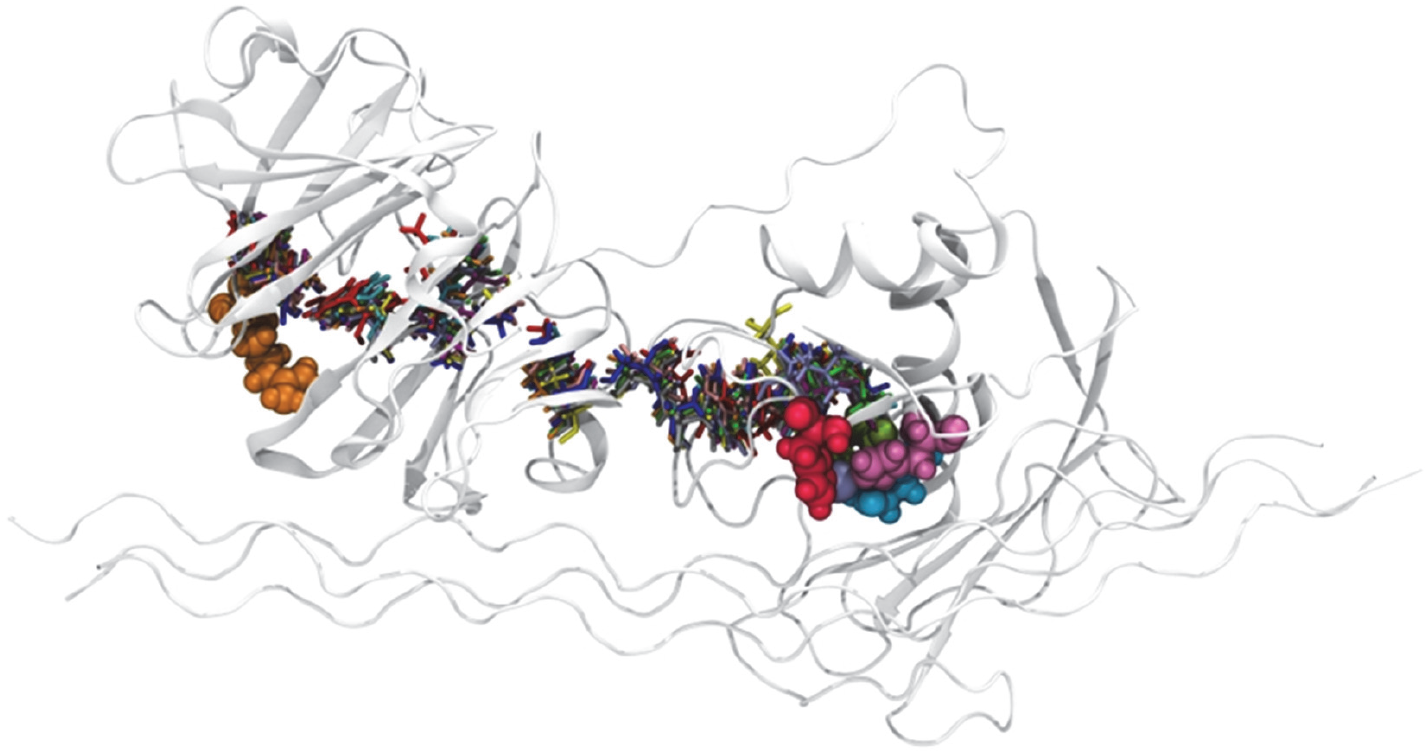
Pathways depicting the top allosteric coupling intensity values from MMP1 with unrestrained collagen. Singular pathways are represented by color.

**Figure S6.**
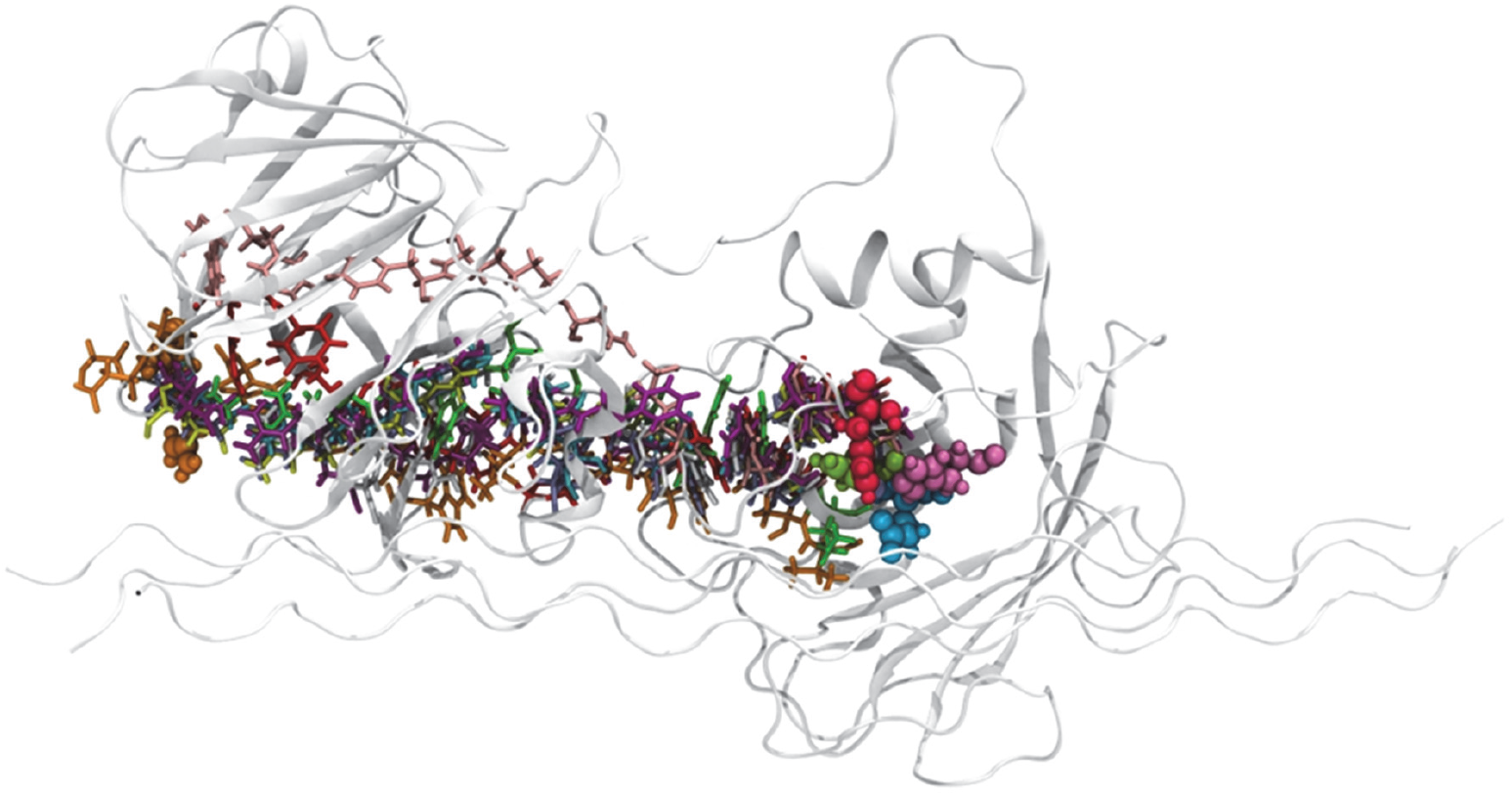
Pathways depicting the top allosteric coupling intensity values from MMP1 with 1000 kJ/mol on the collagen backbone alpha-carbons. Singular pathways are represented by color.

**Figure S7.**
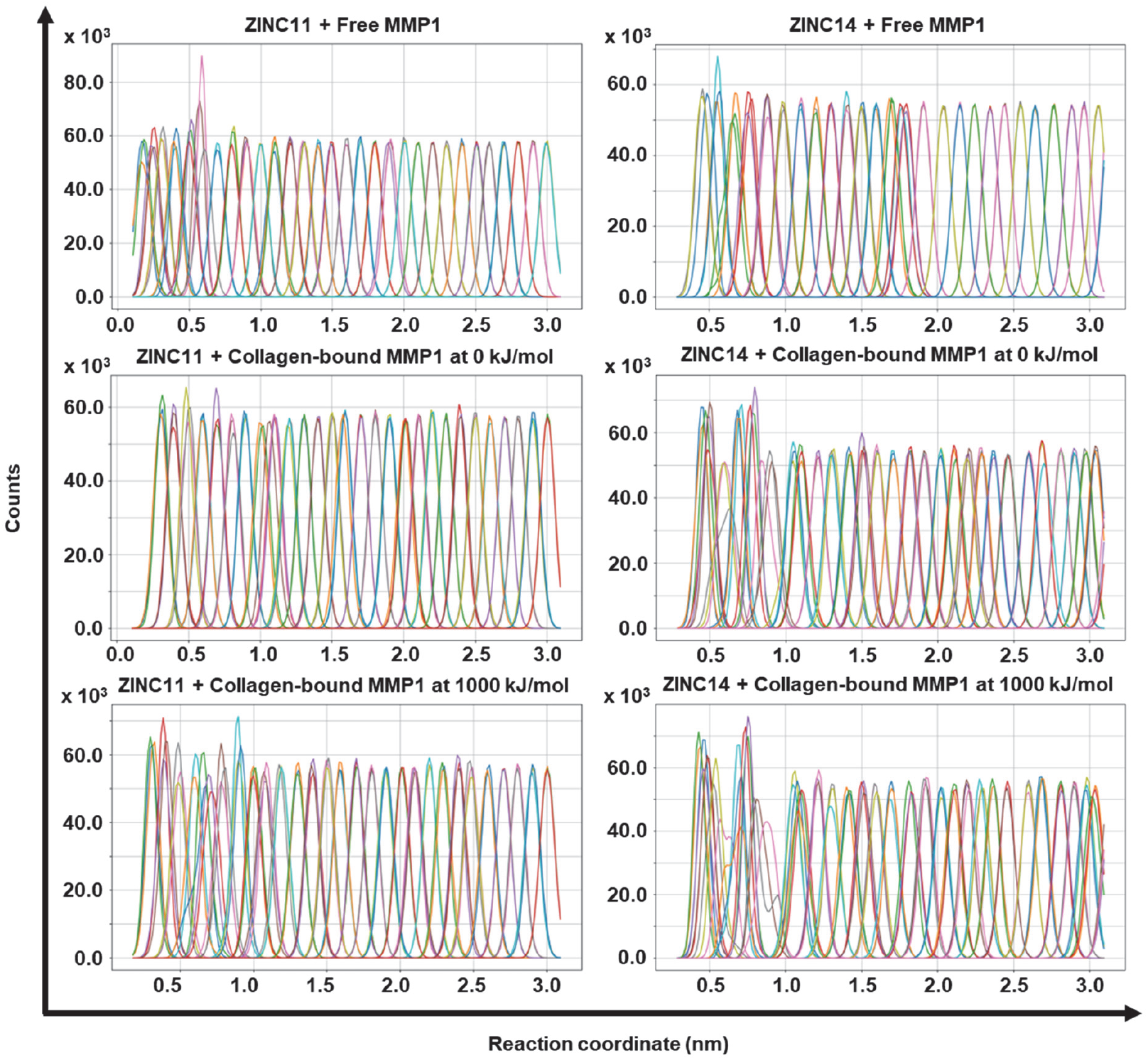
Individual PMF simulations were performed in triplicate for 40 ns every 0.1 nm along the reaction coordinate. The histograms show a well-sampled reaction coordinate, with sufficient overlap between adjacent simulations. Due to the small molecule’s proximity relative to the protein, the taller peaks at the start of the reaction coordinate are due to the small molecule having fewer degrees of freedom.

**Figure S8.**
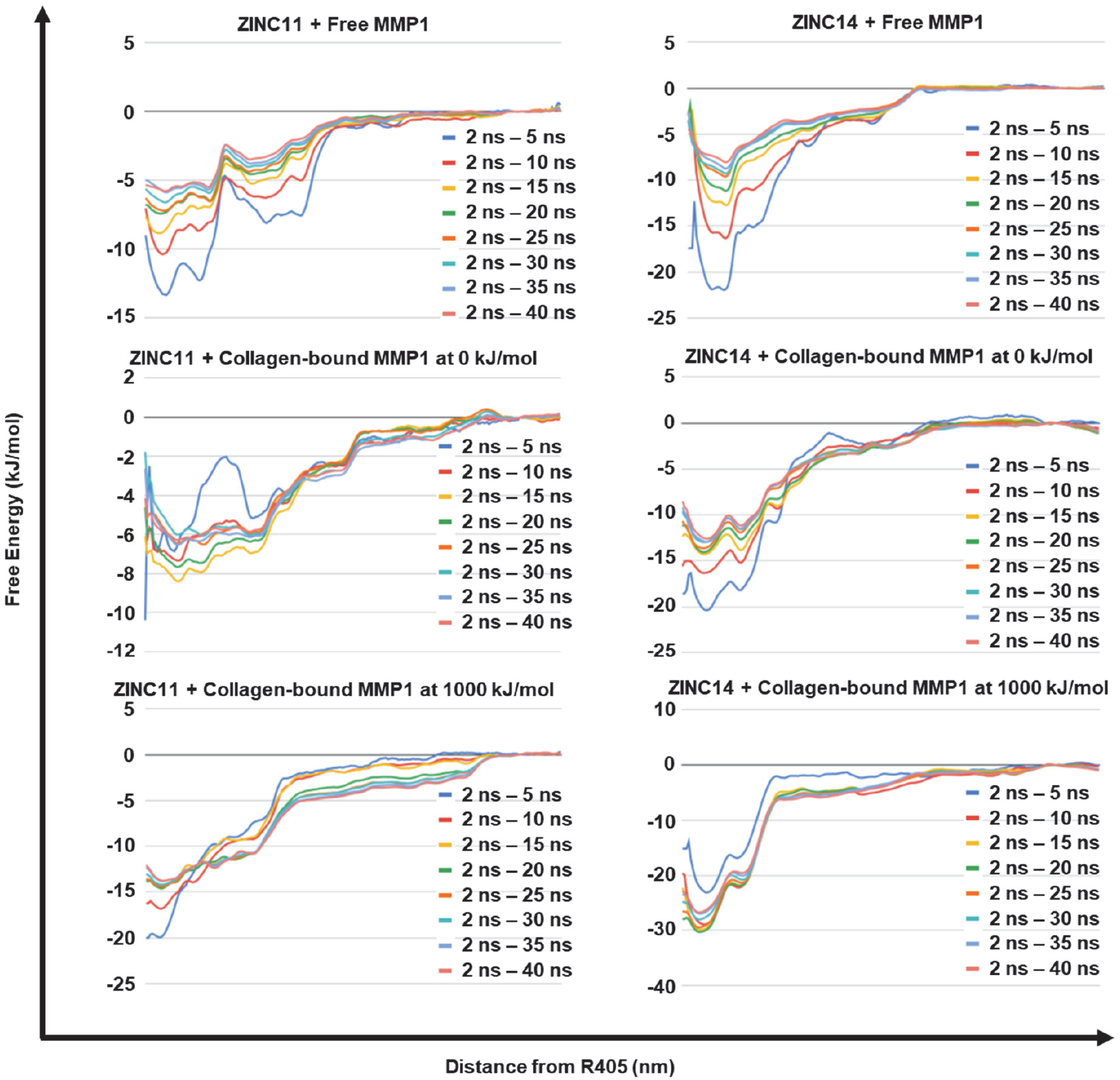
PMF calculations were checked for convergence by systematically adding additional sampling time steps until the free energy plot stopped adjusting. The first sample duration was 2 ns—5 ns, with the first 2 ns disregarded to ensure the system had fully relaxed. Additional 5 ns simulation time frames were then added.

## References

1. H. Nagase, R. Visse, G. Murphy, Structure and function of matrix metalloproteinases and TIMPs. Cardiovascular Research 69, 562–573 (2006).

2. S. Ricard-Blum, The collagen family. Cold Spring Harbor Perspectives in Biology 3, a004978 (2011).

3. K. Tarnutzer, D. Siva Sankar, J. Dengjel, C. Y. Ewald, Collagen constitutes about 12% in females and 17% in males of the total protein in mice. Scientific Reports 13, 4490 (2023).

4. P. Panwar et al., Changes in structural-mechanical properties and degradability of collagen during aging-associated modifications. Journal of Biological Chemistry 290, 23291–23306 (2015).

5. L. Chung et al., Collagenase unwinds triple-helical collagen prior to peptide bond hydrolysis. The EMBO Journal 23, 3020–3030 (2004).

6. L. Kumar et al., Allosteric communications between domains modulate the activity of matrix metalloprotease-1. Biophysical Journal 119, 360–374 (2020).

7. L. Kumar et al., Activity-dependent interdomain dynamics of matrix metalloprotease-1 on fibrin. Scientific Reports 10, 20615 (2020).

8. S. Kamboj et al., Identification of allosteric fingerprints of alpha-synuclein aggregates in matrix metalloprotease-1 and substrate-specific virtual screening with single molecule insights. Scientific Reports 12, 5764 (2022).

9. S. Kamboj et al., Identification of Substrate-specific Allosteric Fingerprints in Matrix Metalloprotease-1 on Amyloid-beta Peptide Aggregates and Drug Screening With Single Molecule Insights. Research Square (2022).

10. D. Rodríguez, C. J. Morrison, C. M. Overall, Matrix metalloproteinases: what do they not do? New substrates and biological roles identified by murine models and proteomics. Biochimica et Biophysica Acta (BBA)-Molecular Cell Research 1803, 39–54 (2010).

11. B. I. Ratnikov et al., Basis for substrate recognition and distinction by matrix metalloproteinases. Proceedings of the National Academy of Sciences 111, E4148–E4155 (2014).

12. D. E. Koshland Jr, G. Némethy, D. Filmer, Comparison of experimental binding data and theoretical models in proteins containing subunits. Biochemistry 5, 365–385 (1966).

13. D. E. Koshland Jr, Application of a theory of enzyme specificity to protein synthesis. Proceedings of the National Academy of Sciences 44, 98–104 (1958).

14. D. E. Koshland Jr, The key–lock theory and the induced fit theory. Angewandte Chemie International Edition in English 33, 2375–2378 (1995).

15. J.-P. Changeux, Allostery and the Monod-Wyman-Changeux model after 50 years. Annual review of biophysics 41, 103–133 (2012).

16. J.-P. Changeux, S. Edelstein, Conformational selection or induced fit? 50 years of debate resolved. F1000 biology reports 3 (2011).

17. D. D. Boehr, R. Nussinov, P. E. Wright, The role of dynamic conformational ensembles in biomolecular recognition. Nature chemical biology 5, 789–796 (2009).

18. B. Ma, M. Shatsky, H. J. Wolfson, R. Nussinov, Multiple diverse ligands binding at a single protein site: a matter of pre-existing populations. Protein science 11, 184–197 (2002).

19. S. Kumar, B. Ma, C.-J. Tsai, N. Sinha, R. Nussinov, Folding and binding cascades: dynamic landscapes and population shifts. Protein science 9, 10–19 (2000).

20. C.-J. Tsai, B. Ma, R. Nussinov, Folding and binding cascades: shifts in energy landscapes. Proceedings of the National Academy of Sciences 96, 9970–9972 (1999).

21. C.-J. Tsai, S. Kumar, B. Ma, R. Nussinov, Folding funnels, binding funnels, and protein function. Protein Science 8, 1181–1190 (1999).

22. B. Ma, S. Kumar, C.-J. Tsai, R. Nussinov, Folding funnels and binding mechanisms. Protein engineering 12, 713–720 (1999).

23. G. G. Hammes, Y.-C. Chang, T. G. Oas, Conformational selection or induced fit: a flux description of reaction mechanism. Proceedings of the National Academy of Sciences 106, 13737–13741 (2009).

24. X. Deupi, B. K. Kobilka, Energy landscapes as a tool to integrate GPCR structure, dynamics, and function. Physiology 25, 293–303 (2010).

25. A. S. Raman, K. I. White, R. Ranganathan, Origins of allostery and evolvability in proteins: a case study. Cell 166, 468–480 (2016).

26. V. H. Salinas, R. Ranganathan, Coevolution-based inference of amino acid interactions underlying protein function. Elife 7, e34300 (2018).

27. G. M. Süel, S. W. Lockless, M. A. Wall, R. Ranganathan, Evolutionarily conserved networks of residues mediate allosteric communication in proteins. Nature structural biology 10, 59–69 (2003).

28. A. Goncearenco et al., SPACER: server for predicting allosteric communication and effects of regulation. Nucleic acids research 41, W266–W272 (2013).

29. J. Monod, J. Wyman, J.-P. Changeux, On the nature of allosteric transitions: A plausible model. Journal of Molecular Biology 12, 88–118 (1965).

30. A. Cooper, D. Dryden, Allostery without conformational change: a plausible model. European Biophysics Journal 11, 103–109 (1984).

31. C. J. Tsai, S. Kumar, B. Ma, R. Nussinov, Folding funnels, binding funnels, and protein function. Protein Science 8, 1181–1190 (1999).

32. S. W. Lockless, R. Ranganathan, Evolutionarily conserved pathways of energetic connectivity in protein families. Science 286, 295–299 (1999).

33. K. Gunasekaran, B. Ma, R. Nussinov, Is allostery an intrinsic property of all dynamic proteins? Proteins: Structure, Function, and Bioinformatics 57, 433–443 (2004).

34. C.-J. Tsai, R. Nussinov, A unified view of “how allostery works”. PLoS Comput Biol 10, e1003394 (2014).

35. R. Kalescky, H. Zhou, J. Liu, P. Tao, Rigid residue scan simulations systematically reveal residue entropic roles in protein allostery. PLoS computational biology 12, e1004893 (2016).

36. M. D. Daily, J. J. Gray, Allosteric communication occurs via networks of tertiary and quaternary motions in proteins. PLoS computational biology 5, e1000293 (2009).

37. E. Guarnera, I. N. Berezovsky, Toward Comprehensive Allosteric Control over Protein Activity. Structure 27, 866-878. e861 (2019).

38. B. Ma, C.-J. Tsai, T. Haliloğlu, R. Nussinov, Dynamic allostery: linkers are not merely flexible. Structure 19, 907–917 (2011).

39. I. Kass, A. Horovitz, Mapping pathways of allosteric communication in GroEL by analysis of correlated mutations. Proteins: Structure, Function, and Bioinformatics 48, 611–617 (2002).

40. J. O. Wrabl et al., The role of protein conformational fluctuations in allostery, function, and evolution. Biophysical chemistry 159, 129–141 (2011).

41. S. W. Manka et al., Structural insights into triple-helical collagen cleavage by matrix metalloproteinase 1. Proceedings of the National Academy of Sciences 109, 12461–12466 (2012).

42. H. Topol, H. Demirkoparan, T. J. Pence, Fibrillar collagen: a review of the mechanical modeling of strain-mediated enzymatic turnover. Applied Mechanics Reviews 73, 050802 (2021).

43. A. P. Bhole et al., Mechanical strain enhances survivability of collagen micronetworks in the presence of collagenase: implications for load-bearing matrix growth and stability. Philosophical Transactions of the Royal Society A: Mathematical, Physical and Engineering Sciences 367, 3339–3362 (2009).

44. R. J. Camp et al., Molecular mechanochemistry: low force switch slows enzymatic cleavage of human type I collagen monomer. Journal of the American chemical society 133, 4073–4078 (2011).

45. B. P. Flynn et al., Mechanical strain stabilizes reconstituted collagen fibrils against enzymatic degradation by mammalian collagenase matrix metalloproteinase 8 (MMP-8). PloS one 5, e12337 (2010).

46. J. C. Lotz, T. Hadi, C. Bratton, K. M. Reiser, A. H. Hsieh, Anulus fibrosus tension inhibits degenerative structural changes in lamellar collagen. European Spine Journal 17, 1149-1159 (2008).

47. J. C. Ellsmere, R. A. Khanna, J. M. Lee, Mechanical loading of bovine pericardium accelerates enzymatic degradation. Biomaterials 20, 1143–1150 (1999).

48. A. S. Adhikari, J. Chai, A. R. Dunn, Mechanical load induces a 100-fold increase in the rate of collagen proteolysis by MMP-1. Biophysical Journal 100, 513a (2011).

49. A. S. Adhikari, E. Glassey, A. R. Dunn, Conformational dynamics accompanying the proteolytic degradation of trimeric collagen I by collagenases. Journal of the American Chemical Society 134, 13259–13265 (2012).

50. E. Yi et al., Mechanical forces accelerate collagen digestion by bacterial collagenase in lung tissue strips. Frontiers in physiology 7, 287 (2016).

51. S. Ghazanfari, A. Driessen-Mol, C. Bouten, F. Baaijens, Modulation of collagen fiber orientation by strain-controlled enzymatic degradation. Acta biomaterialia 35, 118–126 (2016).

52. C. Huang, I. Yannas, Mechanochemical studies of enzymatic degradation of insoluble collagen fibers. Journal of biomedical materials research 11, 137–154 (1977).

53. X. Wang, X. Shen, X. Li, C. M. Agrawal, Age-related changes in the collagen network and toughness of bone. Bone 31, 1–7 (2002).

54. A. Dittmore et al., Internal strain drives spontaneous periodic buckling in collagen and regulates remodeling. Proceedings of the National Academy of Sciences 113, 8436–8441 (2016).

55. I. Bertini et al., Structural basis for matrix metalloproteinase 1-catalyzed collagenolysis. Journal of the American Chemical Society 134, 2100–2110 (2012).

56. L. Cerofolini et al., Examination of matrix metalloproteinase-1 in solution: a preference for the pre-collagenolysis state. Journal of Biological Chemistry 288, 30659–30671 (2013).

57. L. H. Arnold et al., The interface between catalytic and hemopexin domains in matrix metalloproteinase-1 conceals a collagen binding exosite. Journal of Biological Chemistry 286, 45073–45082 (2011).

58. T. G. Karabencheva-Christova, C. Z. Christov, G. B. Fields, Conformational dynamics of matrix metalloproteinase-1· triple-helical peptide complexes. The Journal of Physical Chemistry B 122, 5316–5326 (2017).

59. I. Bertini et al., Interdomain flexibility in full-length matrix metalloproteinase-1 (MMP-1). Journal of Biological Chemistry 284, 12821–12828 (2009).

60. W. Singh, G. B. Fields, C. Z. Christov, T. G. Karabencheva-Christova, Importance of the linker region in matrix metalloproteinase-1 domain interactions. RSC advances 6, 23223–23232 (2016).

61. H. Piccard, P. E. Van den Steen, G. Opdenakker, Hemopexin domains as multifunctional liganding modules in matrix metalloproteinases and other proteins. Journal of Leucocyte Biology 81, 870–892 (2007).

62. U. Eckhard et al., Active site specificity profiling of the matrix metalloproteinase family: Proteomic identification of 4300 cleavage sites by nine MMPs explored with structural and synthetic peptide cleavage analyses. Matrix biology 49, 37–60 (2016).

63. J. Wang et al., Mapping allosteric communications within individual proteins. Nature Communications 11, 3862 (2020).

64. S. K. Sarkar, B. Marmer, G. Goldberg, K. C. Neuman, Single-molecule tracking of collagenase on native type I collagen fibrils reveals degradation mechanism. Current Biology 22, 1047–1056 (2012).

65. M. F. Browner, W. W. Smith, A. L. Castelhano, Matrilysin-inhibitor complexes: common themes among metalloproteases. Biochemistry 34, 6602–6610 (1995).

66. W. Kester, B. Matthews, Crystallographic study of the binding of dipeptide inhibitors to thermolysin: implications for the mechanism of catalysis. Biochemistry 16, 2506–2516 (1977).

67. S. Manzetti, D. R. McCulloch, A. C. Herington, D. van der Spoel, Modeling of enzyme– substrate complexes for the metalloproteases MMP-3, ADAM-9 and ADAM-10. Journal of computer-aided molecular design 17, 551–565 (2003).

68. P. S. Nerenberg, R. Salsas-Escat, C. M. Stultz, Do collagenases unwind triple-helical collagen before peptide bond hydrolysis? Reinterpreting experimental observations with mathematical models. Proteins: Structure, Function, and Bioinformatics 70, 1154–1161 (2008).

69. B. I. Tingle et al., ZINC-22─ A free multi-billion-scale database of tangible compounds for ligand discovery. Journal of chemical information and modeling 63, 1166–1176 (2023).

70. C. A. Lipinski, Lead-and drug-like compounds: the rule-of-five revolution. Drug discovery today: Technologies 1, 337–341 (2004).

71. T. F. Vieira, S. F. Sousa, Comparing AutoDock and Vina in ligand/decoy discrimination for virtual screening. Applied Sciences 9, 4538 (2019).

72. J. Eberhardt, D. Santos-Martins, A. F. Tillack, S. Forli, AutoDock Vina 1.2. 0: New docking methods, expanded force field, and python bindings. Journal of chemical information and modeling 61, 3891–3898 (2021).

73. M. Karplus, Role of conformation transitions in adenylate kinase. Proceedings of the National Academy of Sciences 107, E71–E71 (2010).

74. A. V. Pisliakov, J. Cao, S. C. Kamerlin, A. Warshel, Enzyme millisecond conformational dynamics do not catalyze the chemical step. Proceedings of the National Academy of Sciences 106, 17359–17364 (2009).

75. P. Li, K. M. Merz Jr, MCPB. py: A python based metal center parameter builder. Journal of Chemical Information and Modeling (2016).

76. R. J. Gowers et al. (2019) MDAnalysis: a Python package for the rapid analysis of molecular dynamics simulations. (Los Alamos National Laboratory (LANL), Los Alamos, NM (United States)).

77. N. Michaud-Agrawal, E. J. Denning, T. B. Woolf, O. Beckstein, MDAnalysis: a toolkit for the analysis of molecular dynamics simulations. Journal of computational chemistry 32, 2319–2327 (2011).

78. M. J. Abraham et al., GROMACS: High performance molecular simulations through multi-level parallelism from laptops to supercomputers. SoftwareX 1, 19–25 (2015).

79. L. Kumar et al., Activity-dependent interdomain dynamics of matrix metalloprotease-1 on fibrin. Scientific Reports 10, 1–14 (2020).

80. J. S. Hub, B. L. De Groot, D. Van Der Spoel, g_wham A Free Weighted Histogram Analysis Implementation Including Robust Error and Autocorrelation Estimates. Journal of chemical theory and computation 6, 3713–3720 (2010).

## References

1. S. A. Hollingsworth, R. O. Dror, Molecular dynamics simulation for all. Neuron 99, 1129–1143 (2018).

